# Diet modifies allele-specific phenotypes in *Drosophila* carrying epilepsy-associated *PNPO* variants

**DOI:** 10.1101/2021.07.19.452889

**Authors:** Wanhao Chi, Atulya SR Iyengar, Wenqin Fu, Wei Liu, Abigayle E Berg, Chun-Fang Wu, Xiaoxi Zhuang

## Abstract

Pyridox(am)ine 5’-phosphate oxidase (PNPO) catalyzes the rate-limiting step in the synthesis of pyridoxal 5’-phosphate (PLP), the active form of vitamin B6 required for the synthesis of neurotransmitters GABA and monoamines. Pathogenic variants in *PNPO* have been repeatedly identified in patients with neonatal epileptic encephalopathy and early-onset epilepsy. These patients often exhibit different types of seizures and variable comorbidities, including developmental impairment and intellectual disability. It is unclear how seizure types and associated comorbidities are linked to specific *PNPO* alleles and to what degree diet can modify their expression. Furthermore, the molecular characteristics of *PNPO* variants have not been examined in model systems. Using CRISPR/Cas9, we generated four knock-in *Drosophila* alleles, *h^WT^, h^R116Q^, h^D33V^*, and *h^R95H^*, in which the endogenous *Drosophila PNPO* (*sugarlethal*) was replaced by wild-type human *PNPO* cDNA and epilepsy-associated variants corresponding to R116Q, D33V, and R95H, respectively. We examined these knock-in flies at the molecular, circuitry, and behavioral levels. Collectively, we found a wide range of phenotypes in an allele- and diet-dependent manner. Specifically, the D33V mutation reduces the mRNA level, R95H reduces the protein stability, and R116Q alters the protein localization of PNPO in the brain. D33V and R95H mutations lead to partial and complete lethality during development, respectively and R116Q and D33V mutations shorten lifespan. At the behavioral level, *h^D33V^/h^R95H^* trans-heterozygous flies are hypoactive on all tested diets whereas *h^R116Q^* flies show diet-dependent locomotor activities. At the circuitry level, *h^D33V^* homozygotes show rhythmic burst firing and *h^D33V^/h^R95H^* trans-heterozygotes exhibit spontaneous seizure discharges. In *h^R95H^* homozygotes rescued with PLP supplementation, we uncovered that PLP deficiency abolishes development and causes extreme seizures in adults. Lastly, genetic and electrophysiological analyses demonstrate that *h^WT^/h^R95H^* heterozygous flies are prone to seizures due to a dominant-negative effect of h^R95H^ on h^WT^, highlighting the possibility that human R95H carriers may also be susceptible to epilepsy. Together, this study demonstrates that human *PNPO* variants interact with diet to contribute to phenotypic variations; and that the knock-in *Drosophila* model offers a powerful approach to systematically examine clinical manifestations and the underlying mechanisms of human PNPO deficiency.

## Introduction

Pyridox(am)ine 5’-phosphate oxidase (PNPO, EC 1.4.3.5) is a rate-limiting enzyme in the synthesis of vitamin B6 (VB6) [1]. Mutations in PNPO can cause neonatal epileptic encephalopathy, a devastating disease that usually leads to death if untreated [2]. Recently, PNPO mutations have also been reported in patients with infantile spasms and early-onset epilepsy [3,4,5,6], and the *PNPO* gene has been recognized as one of the sixteen epilepsy genes involved in the genetic generalized epilepsy in adults [7]. The increasingly recognized impact of *PNPO* in epilepsy raises the question of how *PNPO* variants are implicated in different types of epilepsy and ages of seizure onset.

VB6 comprises pyridoxine (PN), pyridoxamine (PM), pyridoxal (PL), and their corresponding phosphorylated forms. Among them, PLP is the only active form, which is a cofactor for more than 140 enzymes including those required for the synthesis of neurotransmitters dopamine, serotonin, and GABA [8]. Unlike plants, bacteria, and fungi, humans and other animals cannot synthesize VB6 de novo, instead relying on dietary VB6 which is converted to PLP by PNPO [1]. The amino acid sequence and structure of PNPO are conserved well from *E. coli* to humans [9,10,11]. In mammals, PNPO is highly expressed in the liver, kidney, and brain [12]. The functional PNPO protein is a homodimer and it binds to two molecules of flavin mononucleotide (FMN) as cofactors [13,14,15,16].

To date, more than thirty *PNPO* variants have been identified in neonatal epileptic encephalopathy and other epilepsy patients since the first report in 2005 [2,17,18]. Biochemical studies show that different variants reduce the PNPO enzymatic activity to different levels, ranging from 0% to 83% [2,3,19,20]. Such a wide range of variation can be attributed to the differential effects of different mutations on the catalytic site, FMN binding, and/or protein folding and thermostability [18,21,22]. While these in vitro studies show clear evidence that different PNPO mutations affect enzymatic activity to varying degrees, they do not explain the variation in seizure types, seizure onsets, and comorbidities manifested by PNPO deficient patients carrying the same mutation [3,18]. Furthermore, *PNPO* variants have never been studied *in vivo* for their molecular or functional effects. It remains unknown whether they can also indirectly affect the gene’s function through, for example, affecting the mRNA/protein level or protein localization. These molecular characterizations are presumably crucial for understanding the phenotypic variation from a molecular perspective. Therefore, there is a need to develop in vivo systems to better understand individual *PNPO* variants and their potential phenotypic variations.

Here we generated four *Drosophila* knock-in (KI) strains that carry either a wild-type (WT) human *PNPO* allele (*h^WT^*) or one of three human *PNPO* epilepsy-associated alleles *h^R116Q^, h^D33V^*, and *h^R95H^*. We examined KI flies at the molecular, circuitry, and behavioral levels. In addition to the reported impaired enzymatic activity, we found that each mutant variant confers specific molecular effects: *h^D33V^* decreases mRNA level, *h^R95H^* reduces protein stability, and *h^R116Q^* alters the protein localization of *PNPO* in the brain. Furthermore, we observed a wide range of phenotypes in KI flies, including developmental impairment, behavioral hyper- or hypo-activity, spontaneous seizure discharges or abnormal firing patterns, and shortened lifespan. The phenotypic variation is associated with the known severity of these mutations and our newly characterized molecular effects. We also showed that diet treatments further diversified the phenotypes among alleles. Lastly, we found that *h^R95H^* had a significant dominant-negative (DN) effect and heterozygous flies were prone to seizures upon electroconvulsive stimulation.

## Methods

### Generation of knock-in strains

Four different KI strains were generated using CRISPR/Cas9 technology [23].The *Drosophila PNPO* gene (*sugarlethal*) [10] was replaced by either WT human *PNPO* (*hPNPO*) cDNA or one of three mutant *hPNPO* cDNAs. The WT *hPNPO* cDNA was amplified from the human brain cDNA library (TaKaRa, Cat #637242) [24]. The c.98A >T mutation (corresponding to p.D33V), c.284G > A mutation (corresponding to p.R95H), and c.347G > A mutation (corresponding to p.R116Q) were introduced separately by mutagenesis. The single guide RNAs (sgRNAs) were designed with CRISPR Optimal Target Finder [25] and transcribed *in vitro* as described in the published protocol [26] (Supplementary Table 1). Cas9 messenger RNAs (mRNAs) were transcribed *in vitro* from plasmid MLM3613 (Addgene, plasmid #42251). Donors were various *hPNPO* cDNAs assembled in the pBluescript SK(-) vector. The sgRNAs, Cas9 mRNAs, and donor constructs were injected into embryos from flies with a genotype of *w^1118^/FM7a; Bc/CyO; TM3/TM6B, Hu, Tb* (www.fungene.tech). The introduction of the mutation in KI alleles was confirmed by Sanger sequencing of PCR products amplified with a pair of PCR primers that specifically targeted *hPNPO* cDNA (Supplementary Table 1).

The mutant alleles were initially maintained in heterozygotes with the *TM6B, Hu, Tb* balancer, but KI homozygotes gradually took over heterozygotes in *h^WT^* and *h^R116Q^* breeding bottles.

### *Drosophila* husbandry

Electrophysiology experiments and some behavioral recordings (indicated below) were performed on flies bred on Frankel Brosseau’s (FB) media [27] at the University of Iowa. For all other experiments, flies were generated on standard Cornmeal-Yeast-Molasses (CYM) media from the Fly Kitchen at the University of Chicago. Flies used in all experiments were raised and tested at room temperature (~ 23 °C) in a 12:12 hour light:dark cycle.

### Western blotting

Male flies, 1- to 2-day old, were used. Total protein from thirty male adult heads was extracted and quantified. A total of 50 μg protein from each sample was loaded for SDS-PAGE. Separated proteins were electrophorectically transferred to the PVDF membrane. After blocking, the membrane was incubated with primary antibodies and then the secondary antibodies (Supplementary Table 2). Signals were detected with chemiluminescence (ThermoScientific, Cat #32109).

### RNA extraction, reverse transcription, and quantitative PCR

Total RNA was extracted from thirty male adult heads using the RNA extraction kit (Zymo Research). After removal of genomic DNA using the DNA-free kit (AMBION, Cat #R2030), total RNA was used for reverse transcription (RT) using the SMARTScribe Reverse Transcription Kit (TaKaRa, Cat #639537).

Quantitative PCR (qPCR) was performed using SYBR Green real-time PCR method (Applied biosystems). The Ct values were defined by the default settings in QuantStudio3 (Thermofisher) using a run method of 2 minutes at 95 °C followed by 40 cycles of 15 seconds at 95 °C and 1 minute at 60 °C. Relative gene expression was calculated as 2^ΔCt^, where ΔCt was determined by subtracting the average Ct value from the housekeeping gene ribosomal protein 49 (*rp49*).

Two pairs of primers were designed to target the *hPNPO* KI alleles (N- and C- terminal regions, respectively). A pair of primers were used to amplify *rp49*. Primers were pre-tested for single-product amplification. Primer sequences are shown in Supplementary Table 1.

### Immunohistochemistry staining and confocal imaging

The protocol was adapted from the Flylight protocol [28]. Briefly, brains from 1- to 2-day old flies were dissected in cold S2 media and fixed with 2% paraformaldehyde for 55 minutes. After a brief rinse with 0.5% Triton X-100, brains were blocked with blocking solution (5% normal donkey serum and 0.5% TritonX-100 in phosphate-buffered saline) for 1.5 hours and then were incubated with primary antibodies (Supplementary Table 2). Signals for hPNPO were amplified with the Tyramide SuperBoost kit (Invitrogen, Cat # B40926) by following the manufacturer’s protocol. After incubation with the secondary antibodies and washes, brains were mounted on a double-frosted slide and covered with coverslips. DAPI was added into the wash buffer to stain the nucleus when needed. Images were taken using Leica SP5-II-STED-CW confocal microscope in the Integrated Light Microscopy Core Facility at the University of Chicago and processed in Fiji [29].

### Developmental assay with or without PLP supplementation

Male and female heterozygous flies were bred in normal food vials or bottles with or without PLP supplementation. A cohort of two to three flies per sex was set up in a single vial, or a group of twelve to fifteen flies per sex was set up in a single bottle. F1 flies eclosed within six days from each cross were examined for the Balancer marker.

### Lifespan and survival study

Fifteen to twenty male flies, 1- to 2-day old, were maintained in vials filled with the standard CYM medium (the normal diet condition) or 4% sucrose in 1% agar (the sugar-only condition). When there was PLP supplementation, PLP was added to the vial to the final concertation of 400 *μ*g/ml. Flies were transferred to new vials one to two times a week. Daily survival was recorded.

### Single- or multi-fly behavioral recording

Male flies, 1- to 2-day old, were reared on either control diet, sugar-only diet, or sugar + 400 *μ* g/ml PLP diet for 4 to 6 days. For single-fly recordings, one fly was loaded without anesthesia into a circular arena (diameter = 60 mm) prefilled with 1% agar and was recorded with a camera (FLIR Flea 3) for 3 minutes at 20 fps (frames per second) (Fig. 4A-E). For multi-fly recordings, four flies were loaded without anesthesia into a circular arena (diameter = 28 mm) seated on filter paper (Whatman 1). Flies were recorded with either a webcam (Logitech c920, Fig. 4F-J, 10 minutes at 30 fps), or a camera (Flea3, Fig. 6C-G, 3 minutes at 20 fps). After recording, fly positions in each frame were automatically tracked using IowaFLI Tracker [24,30]. The total distance travelled, average speed, percentage of active time, and speed correlation coefficient (SCC) were further calculated as previously described [24].

### Tethered fly electrophysiology and data analysis

Tethered fly electrophysiological experiments were performed as described previously [24,31]. Briefly, flies were immobilized on ice, affixed onto a tungsten wire between the head and thorax, and allowed 30 minutes for recovery. Flight muscle spikes were recorded by an electrolytically sharpened tungsten electrode inserted into the dorsal longitudinal muscle (DLMa). A sharpened tungsten electrode inserted into the dorsal abdomen served as a reference. Signals were amplified by a Model 1800 AC amplifier (AM systems) and were digitized by a USB 6212 data acquisition card (National Instruments) controlled by a custom LabVIEW 2018 script. Spikes were identified offline by a custom MATLAB script.

The average firing rate was determined by dividing the total spike count during a recording session by the duration (~ 90 seconds). The instantaneous firing frequency was defined as the reciprocal of an inter-spike interval (i.e., ISI^−1^). The instantaneous coefficient of variation (CV_2_), a measure of firing regularity, was defined between two adjacent inter-spike intervals, *i* and *i*+1, as: 2|ISI^−^_*i*_ – ISI^−1^_*i*+*1*_/(ISI^−1^_*i*_ + ISI^−1^_*i*+*1*_) [32]. Lower CV_2_ values indicate with rhythmic spiking, while high CV_2_ values correspond with irregular spiking. Plots of the ISI^−1^ versus CV_2_ trajectories for a spike train were computed as described previously [33].

Systemic application of nicotinic acetylcholine receptor blocker mecamylamine was performed via a dorsal vessel injection procedure [33,34]. Briefly, mecamylamine (Sigma, Cat #M9020) was dissolved in adult hemolymph-like solution [35] to a final concentration of 100 *μ*M and marked with blue #1 dye (16 mg/ml). A filamented glass microelectrode was filled with 0.33 *μ*l mecamylamine solution and inserted into the dorsal vessel. Positive air pressure pushed the solution into the dorsal vessel which systemically circulated the mecamylamine within seconds.

Electroconvulsive seizure discharges were induced by high-frequency electrical stimulation across the brain delivered by tungsten electrodes inserted in each cornea [31,36]. The stimulation protocol consisted of a 2-s train of 0.1 ms pulses delivered at 200 Hz at a specified voltage (30 - 80 V).

### Statistical analysis

Statistical analysis was performed in Matlab (R2019b, U. Iowa) or R (version 3.6.1, U. Chicago). Details on statistical analyses, including sample sizes, tests performed, and multiple test correction if necessary, are provided within the figure legends or included in the figures.

### Data availability

Raw data and customized scripts are available upon request.

## Results

### Generation of knock-in *Drosophila* strains carrying human PNPO variants identified from epilepsy patients

To functionally and molecularly characterize epilepsy-associated variants of *hPNPO*, we generated four KI *Drosophila* alleles using CRISPR/Cas9, in which the *Drosophila PNPO* gene, *sugarlethal* (*sgll*) [10], was replaced by *hPNPO* cDNAs (Fig. 1A). These four KI alleles were designated as *h^R116Q^*, *h^D33V^, h^R95H^*, and *h^WT^*, respectively. The three mutant alleles, *h^R116Q^*, *h^D33V^, h^R95H^*, were chosen to represent different severities of PNPO deficiency based on *in vitro* biochemical studies of their corresponding mutant proteins (h^R116Q^, h^D33V^, h^R95H^ hereafter). The residual enzymatic activity of h^R116Q^, h^D33V^, h^R95H^ are 80%, 40%, and 20% of h^WT^, respectively [3,19]. The mutation in each allele was confirmed by Sanger sequencing (Fig. 1B). All KI alleles were initially balanced over a *TM6B, Hu, Tb* chromosome (*TM6B* hereafter) to circumvent homozygous lethality.

**Figure 1.**
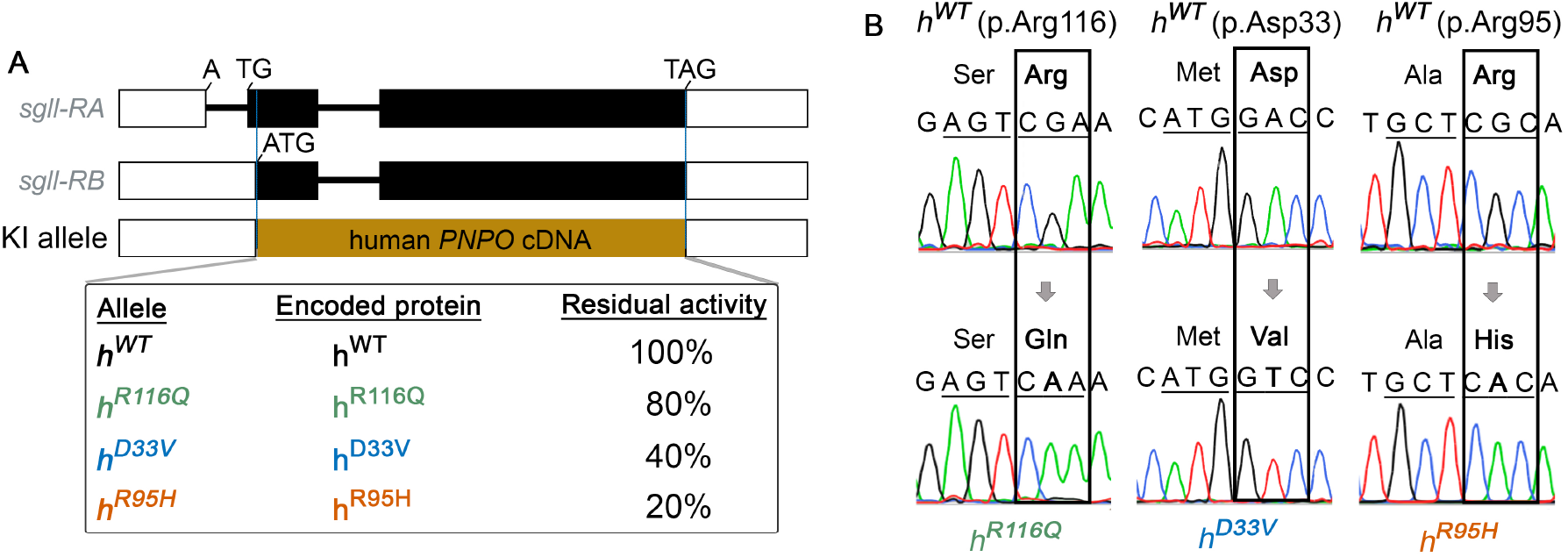
Generation of *Drosophila* knock-in (KI) alleles carrying human *PNPO* (*hPNPO*) epilepsy-associated variants. (**A**) The *Drosophila PNPO* gene, *sgll*, encodes two transcript forms *sgll-RA* and *sgll-RB*. Both of them are expressed; however, *sgll-RB* is globally expressed and has a relatively high expression level in comparison to *sgll-RA* [37].The protein products of these two forms share the C-terminus but differ in the N-terminus, with the sgll-RA form containing nine extra amino acids [38]. Given the first intron in *sgll-RA* could play a regulatory role in gene expression level and/or pattern, *sgll-RB* was replaced by *hPNPO* cDNAs in the KI alleles. Black boxes indicate exons, white boxes indicate UTRs, and black lines indicate introns. KI alleles corresponding to WT, R116Q, D33V, and R95H are designated as *h^WT^*, *h^R116Q^*, *h^D33V^*, and *h^R95H^*, respectively. The residual activities of corresponding *hPNPO* mutant proteins were previously measured biochemically *in vitro* [3,19]. (**B**) Sequencing chromatograms confirmed the presence of the targeted mutation in each KI allele.

### Distinct molecular alterations linked to different *hPNPO* epilepsy-associated variants

Other than reducing the enzymatic activity [2,3,19,20], little is known about the molecular characteristics of *hPNPO* variants. Thus, after establishing the KI lines, we first studied whether these epilepsy-associated variants could affect *hPNPO* expression and localization using homozygous flies (Supplementary Fig. 1). The *h^R95H^* allele was not included here because *h^R95H^* homozygotes were lethal (see below). To include all four KI alleles, we further crossed KI lines with *w^1118^* to generate *KI/sgll^+^* flies and performed western blotting and RT-qPCR using adult heads. We confirmed that the antibody for hPNPO did not recognize *Drosophila* PNPO (Supplementary Fig. 2). Results from western blot demonstrated that the R116Q mutation did not change the hPNPO protein level whereas D33V and R95H significantly reduced it (Fig. 2A-B, Supplementary Fig. 1A-B). Interestingly, multiple bands instead of a single band appeared on the blot. The molecular nature of these bands is unclear. One possibility is that given only the *sgll-RB* form was replaced in the KI allele (Fig. 1A), a chimeric hPNPO protein could also be expressed, which contains the nine extra amino acids from sgll-RA in the front of the hPNPO protein. It is also possible that the multiple bands were due to the proteolysis of the full length of hPNPO, as suggested in a previous study [15].

**Figure 2.**
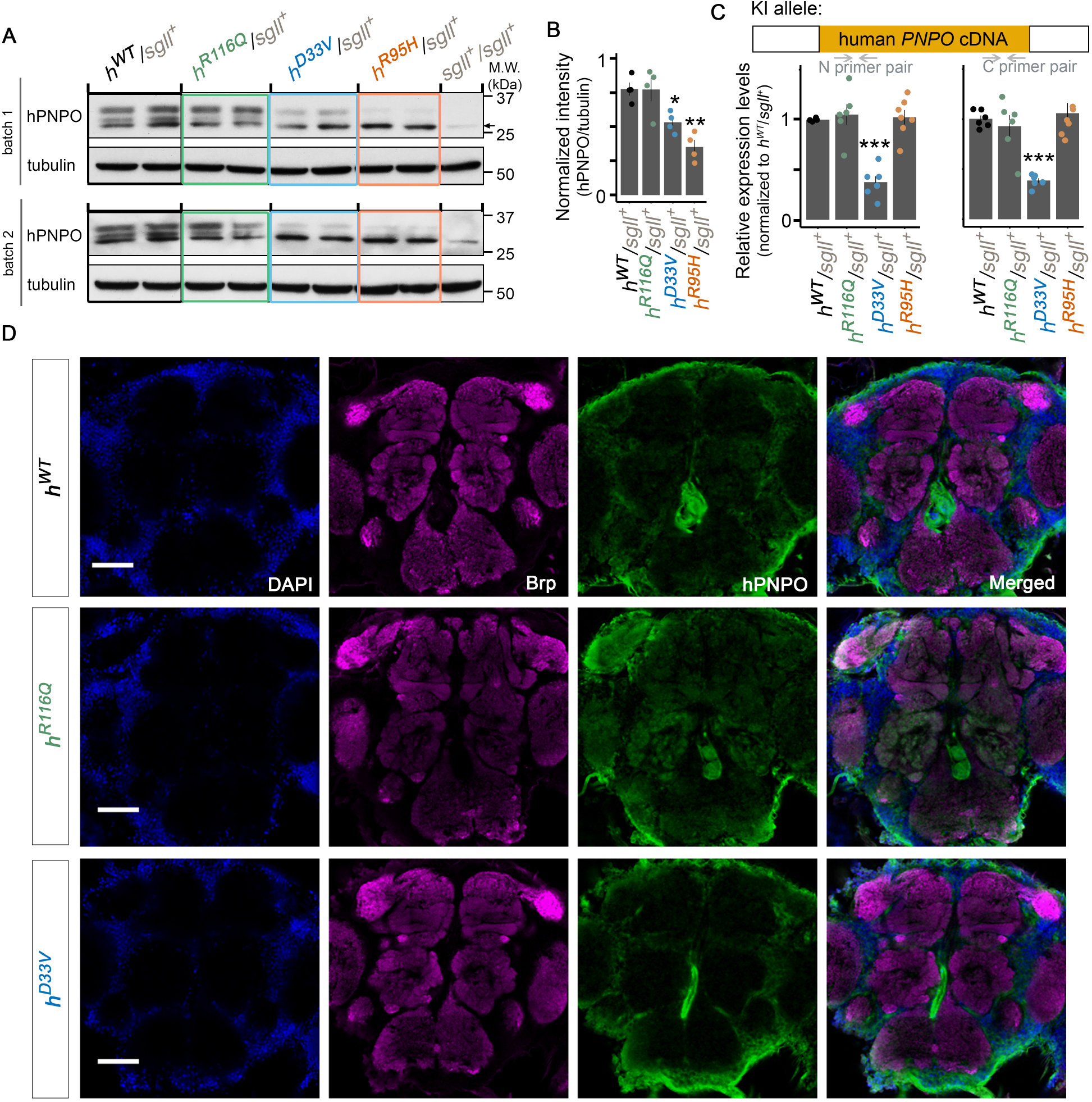
Molecular characterization of each KI allele. (**A**) Western blot of hPNPO in adult head homogenate from each KI/*sgll*^+^ line. n = 4 per genotype. In each batch, one sample from *w^1118^* flies (*sgll*^+^/*sgll*^+^) was loaded as the antibody control. Tubulin was the loading control. The arrow indicates a band also seen in w^1118^ flies, which most likely is a non-specific band (Supplementary Fig. 2). (**B**) Quantification of hPNPO (the top band). n = 4. Error bars represent Mean ±SEM. * *P* < 0.05, ** *P* < 0.01. Two-tailed student’s *t*-test with Bonferroni’s correction compared to *h^WT^*/*sgll*^+^. (**C**) Quantification of the mRNA level of *hPNPO* in adult heads by RT-qPCR. Two primer pairs were used to target the N- and C-terminus of *hPNPO* cDNA, respectively. n = 4-7 per primer pair per genotype. Error bars represent Mean ±SEM. *** *P* < 0.001. Two-tailed student’s t-test with Bonferroni’s correction compared to *h^WT^*/*sgll*^+^. (**D**) Immunohistochemistry staining of hPNPO in the adult brain of *h^WT^*, *h^R116Q^*, and *h^D33V^* homozygotes. The *h^R95H^* line was not included due to the lethality of *h^R95H^* homozygotes in the normal diet and the greatly reduced hPNPO protein level in their heads (panel A). Color representation: Blue, DAPI; Magenta, Brp; Green, hPNPO. Scale: 50 *μ*m.

The reduced hPNPO protein level associated with D33V or R95H mutations could be due to changes at the transcriptional or translational level. We therefore examined the mRNA level of *hPNPO* in each genotype using RT-qPCR. Two pairs of primers that specifically targeted the N- or C-terminal regions of *hPNPO* cDNA were used. Results from both primer pairs showed that D33V but not R95H led to a reduced mRNA level (Fig. 2C, Supplementary Fig. 1C).

Next, we examined the localization of hPNPO protein in the adult brain using immunohistochemistry staining with an anti-hPNPO antibody. Anti-Bruchpilot (Brp) antibody was used to visualize synapse-rich neuropils [39]. We found that h^WT^ was ubiquitously expressed in the brain with the strongest staining in the cell body rind (Fig. 2D, Supplementary Fig. 3) [40]. There was little overlap between Brp and hPNPO staining, suggesting that h^WT^ is not enriched in the terminal structures. A similar staining pattern was observed from h^D33V^ brains (see a representative image in Fig. 2D, n = 5). In striking contrast, strong hPNPO staining in the terminal structures were detected in h^R116Q^ brains. The increased staining appeared to occur in all terminal structures in the brain, with the most dramatic change in the antenna lobe (see a representative image in Fig. 2D, n = 7). Since the hPNPO protein level in h^R116Q^ is comparable to that in h^WT^ (Fig. 2A-B), we conclude that the R116Q mutation alters the hPNPO protein localization in the brain.

Taken together, molecular characterization of *hPNPO* variants in KI flies demonstrates that in addition to directly reducing the enzymatic activity as shown in previous studies [2,3,19,20], *hPNPO* variants can also indirectly affect the gene’s function through altering the mRNA and/or protein level or the protein localization in the brain.

### Distinct developmental effects associated with different *hPNPO* variants

All balanced KI lines were viable and fertile. However, no homozygous KI adult flies were observed from *h^R95H^/TM6B* breeding bottles. To rule out the contribution of any off-target mutations, the KI line was backcrossed to *w^1118^* for five generations. No homozygous *h^R95H^* pupae or adult flies were observed after backcross, suggesting that the lethality was most likely due to the R95H mutation in hPNPO.

To systematically study the effects of the R95H mutation as well as the two other mutations on development, we self-crossed each KI line and examined the number of homozygous KI flies in the F1 generation. The ratio of KI homozygous flies in all flies (homozygosity ratio) was further calculated. Consistent with our initial observations, no *h^R95H^* homozygotes were observed (Fig. 3A). A significantly decreased homozygosity ratio was also found from *h^D33V^* self-breeding. By contrast, the homozygosity ratio in *h^R116Q^* was comparable to that in *h^WT^*. Thus, these three mutant alleles have differential effects on development. This conclusion was corroborated by complementation tests (Supplementary Fig. 4).

**Figure 3.**
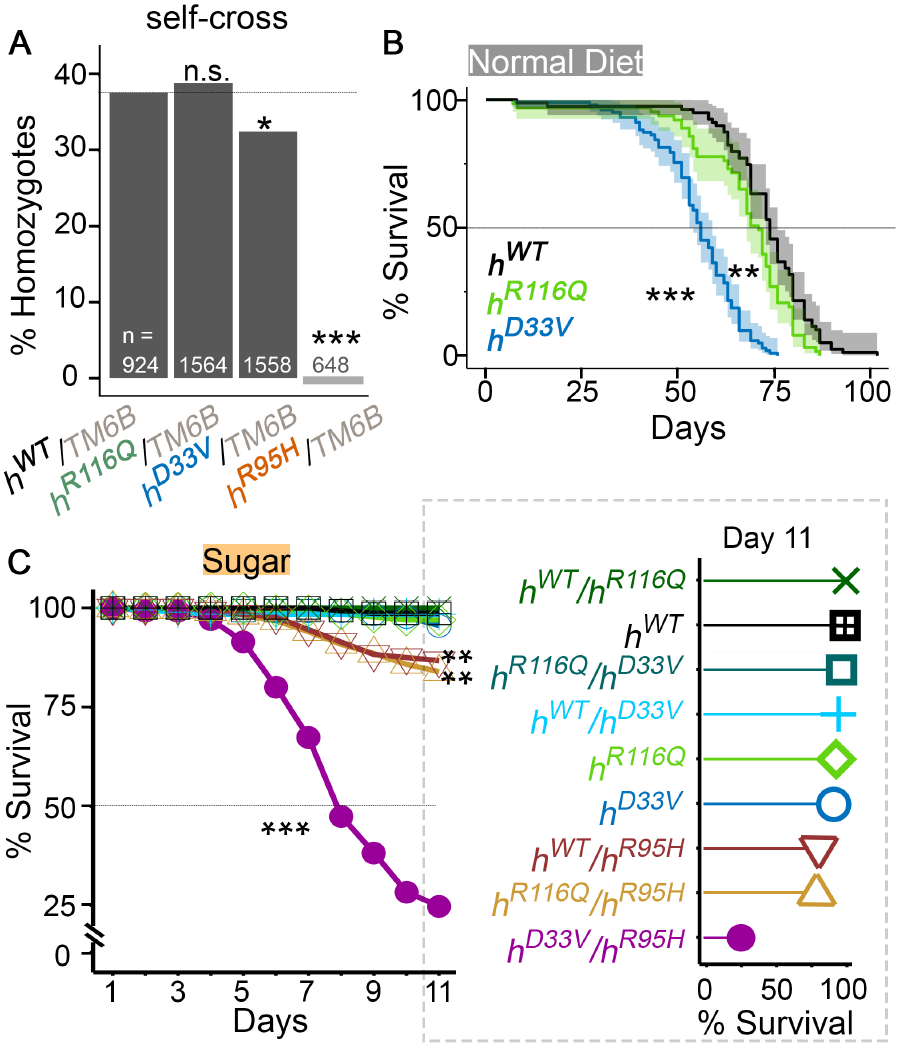
Developmental and survival phenotypes of KI flies from various genotypes. (**A**) Generation of homozygous flies from each KI line. Chi-square test of homogeneity compared to *h^WT^/TM6B* (the gray line). (**B**) Survival analysis of flies on the normal diet. n = 63-102. Log-rank test compared to *h^WT^*. (**C**) Survival of flies from various genotypes on the sugar-only diet. n = 102-218, Log-rank test compared to *h^WT^*. Survival of each genotype on day 11 is also shown in the lollipop plot on the right. n.s. *P* > 0.05, * *P* < 0.05, ** *P* < 0.01, *** *P* < 0.001.

### Allele-dependent diet modifications of lifespan in KI flies

PLP is involved in a variety of biological processes [8], yet the potential cumulative effect of chronic PNPO deficiency has never been studied. We examined whether *h^R116Q^* and *h^D33V^* could affect the survival of adult flies on the normal diet. Survival data showed that *h^R116Q^* had a slightly shortened lifespan compared to *h^WT^* (median: 71 vs. 74 days for *h^R116Q^* and *h^WT^*, respectively. *P* < 0.01. Fig. 3B). In comparison, the lifespan of *h^D33V^* was dramatically shorter (median: 56 days. *P* < 0.001). The lethality of *h^R116Q^* and *h^D33V^* flies correlated well with the residual enzymatic activity of *h^R116Q^* and *h^D33V^* measured by *in vitro* studies [3,19]. Together, these studies demonstrate that even mild PNPO deficiency can have long-term deleterious effects, even in the presence of dietary VB6.

Previously we reported that PNPO deficient flies (*sgll^95^*) had a shortened lifespan on the sugar-only diet (i.e., VB6-devoid diet) [24], suggesting that sugar-only diet is useful for exacerbating PNPO deficiency. We thus generated homozygous (two same KI alleles), heterozygous (one *h^WT^* allele and one mutant allele) and trans-heterozygous (two mutant alleles) flies using four KI alleles and examined their survival on sugar. We found that the most dramatic lethal phenotype among these nine genotypes was from *h^D33V^/h^R95H^* flies (Fig. 3C), of which approximately 75% died by day 11. The significant lethality in *h^D33V^/h^R95H^* flies was not surprising since, based on in vitro residual enzymatic activity studies, *h^D33V^/h^R95H^* has the most severe PNPO deficiency. We also observed lethality from *h^R116Q^/h^R95H^* and *h^WT^/h^R95H^* flies; about 15% of them died by day 11 (Fig. 3C), suggesting that *h^R95H^* is indeed the most severe allele among all three mutant alleles. The fact that *h^WT^/h^R95H^* showed lethality suggests that *h^R95H^* may cause haploinsufficiency or have a DN effect, which is unexpected because PNPO has been considered as autosomal recessive in heritance (https://omim.org). These two possibilities were further studied (see below).

Overall, these studies demonstrate that both *PNPO* variants and dietary conditions can affect survival of KI flies.

### Allele-dependent diet modifications of behaviors of KI flies

The *h^R116Q^* and *h^D33V^* homozygous flies did not exhibit noticeable behavioral deficits when reared on the normal diet (the CYM media). Consistent with this observation, mutant homozygous flies traveled a similar total distance with comparable speed, percentage of active time, and SCC compared to *h^WT^* homozygotes (Supplementary Fig. 5A-E). However, when eclosed flies were reared on sugar for 4-6 days (Fig. 4A), *h^R116Q^* homozygotes exhibited hyperactivity: they traveled farther than *h^WT^* and had an increased average speed (Fig. 4B-E). The increased average speed and total travelled distance were not observed in *h^D33V^* homozygotes in the same breeding and testing conditions (Fig. 4B-E). Interestingly, when files were bred on the other standard diet, the FB media, in which yeast extract and non-fat dry milk were used to replace the dried yeast in the CYM media [27], both *h^R116Q^* and *h^D33V^* homozygous flies exhibited hyperactivity (Supplementary Fig. 5F-J). The difference maintained when eclosed flies were further reared on sugar for 4-6 days (Fig. 4F-J). Therefore, diet plays a significant role in modifying the behaviors of *h^R116Q^* and *h^D33V^* homozygous flies.

**Figure 4.**
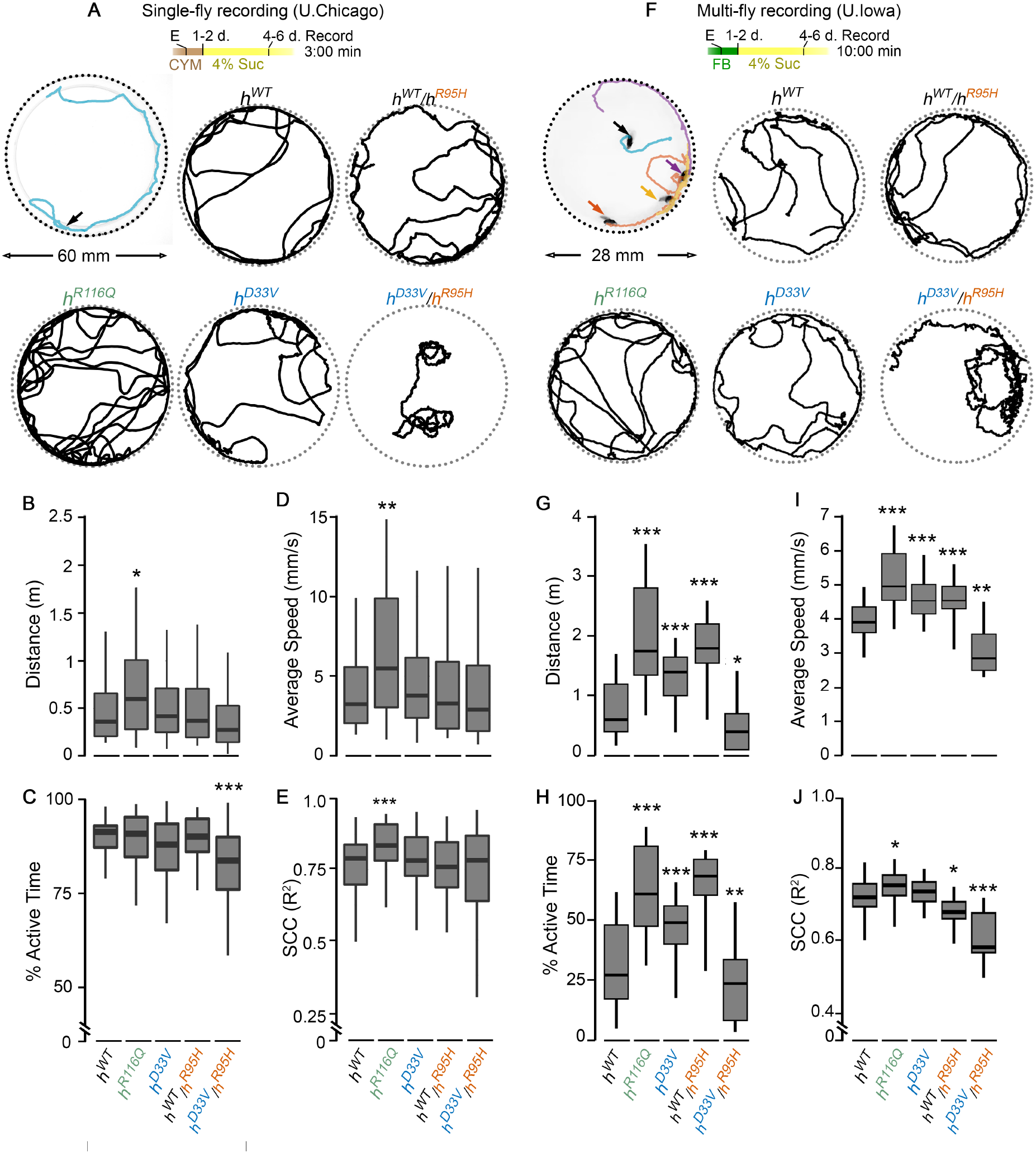
Behavioral analyses of KI flies in an open-field arena. (**A**,**F**) Breeding and testing conditions and representative tracks from each genotype. Flies were eclosed (E) on either CYM or FB diet, and then transferred to the sugar-only diet (4% Suc). (**B-E**) Total distance traveled, percentage of active time, average speed, and speed correlation coefficient (SCC, see Methods for calculation details) of flies from various genotypes generated and tested in condition A. n = 50-75 flies. (**G-J**) Total distance traveled, percentage of active time, average speed, and SCC of flies from various genotypes generated and tested in condition F. n = 28-44 flies. * *P* < 0.05, ** *P* < 0.01, *** *P* < 0.001. Two-tailed student’s *t*-test with Bonferroni’s correction compared to *h^WT^*.

Since we observed lethality in *h^WT^/h^R95H^* and *h^D33V^/h^R95H^* flies when they were reared on sugar (Fig. 3C), we further analyzed their behavior under different breeding and testing dietary conditions. When bred on the CYM media, *h^WT^/h^R95H^* flies behaved similarly to *h^WT^* homozygotes in both sugar-only and CYM testing conditions (Fig. 4B-E, Supplementary Fig. 5B-E). The *h^D33V^/h^R95H^* flies, however, were less active in both testing conditions and some even exhibited tortious walking paths when reared on sugar (see a representative track in Fig. 4A) and, consequently, had low SCCs. The average SCC of *h^D33V^/h^R95H^* flies is comparable to that of *h^WT^* though (Fig. 4E). The fact that the *h^D33V^/h^R95H^* flies exhibit behavioral deficits even on the normal diet suggests that their residual PNPO enzymatic activity is insufficient to convert dietary VB6 to PLP to maintain normal behaviors.

When bred on the FB media, *h^WT^/h^R95H^* flies were more active than *h^WT^* homozygotes in both sugar-only and FB testing conditions (Fig. 4G-J, Supplementary Fig. 5G-J), which resembles *h^R116Q^* and *h^D33V^* homozygotes in the same conditions. The different behaviors of *h^WT^/h^R95H^*, *h^R116Q^*, or *h^D33V^* flies on different media are likely due to the mild PNPO deficiency in them and its interaction with different diets. Consistent with this notion, the *h^D33V^/h^R95H^* flies on the FB media exhibited hypoactivity and tortious walking paths (Fig. 4G-J, Supplementary Video 1), as they were on the CYM media (Fig. 4A-E).

Taken together, these studies demonstrate that diet plays a significant role in the locomotor behaviors of PNPO deficient flies, especially when flies have mild PNPO deficiency.

### Spontaneous seizures in KI flies with severe PNPO deficiency

The spectrum of behavioral phenotypes associated with KI lines prompted us to examine motor unit activity patterns and identify spontaneous seizure-associated spike discharges in the respective lines. We utilized a tethered fly preparation (Fig. 5A) to monitor DLM flight muscle activity in intact, behaving flies. During flight, these muscles power the “down-stroke” of the wing, and the isometric nature of their contractions facilitates prolonged recordings with minimal muscle damage. Importantly, the innervating motor neuron (DLMn) serves an output for several motor programs (e.g., flight, courtship song, and grooming activity) [41,42,43,44]. Several studies have utilized aberrant spiking in this motor unit to monitor spontaneous [45] and evoked [36] seizure activity, including our previous analysis of the original *sgll^95^* mutant [24]. We found *h^WT^* homozygotes reared on sugar displayed occasional grooming-related spiking (Fig. 5A, Supplementary Video 2), largely consistent with activity patterns in other control flies [33]. In the *h^R116Q^* and *h^D33V^* mutants reared on the same diet, we mostly observed similar grooming associated activity patterns, with a marginal increase in the average firing rate in *h^D33V^* flies compared to *h^WT^* controls (Fig. 5B). The increased average firing in *h^D33V^* flies was due to a small subset of *h^D33V^* individuals (3/12) showing a peculiar, highly-stereotypic, pattern of repeated DLM spike bursts (Fig. 5A, Supplementary Video 2). This bursting did not have clear behavioral correlates and was distinct from previously reported spontaneous seizure activity in *sgll^95^* mutants [24].

**Figure 5.**
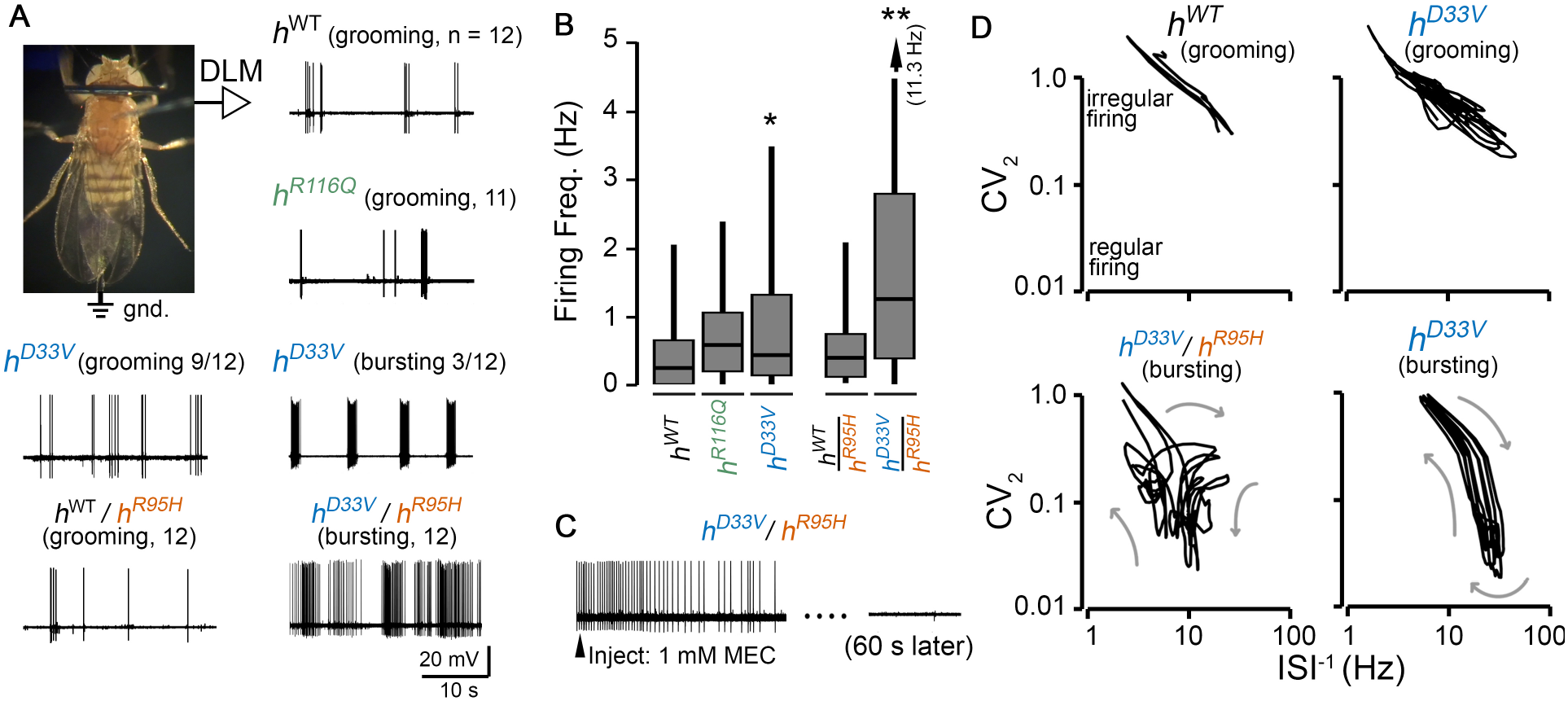
Electrophysiological correlates of seizures-related activity in KI flies. (**A**) Tethered fly preparation (see Methods for details) and representative traces of spontaneous activity in DLM flight muscle spiking flies from each genotype reared on the sugar-only diet. Firing in *h^WT^*, *h^R116Q^*, *h^D33V^* (left trace), and *h^WT^/h^R95H^* are related to grooming behavior. Abnormal rhythmic spike bursts are occasionally observed in *h^D33V^* (right trace, 3/12 individuals), while aberrant spike discharges are observed in *h^D33V^/h^R95H^* flies. (**B**) Average DLM firing rate in flies from each genotype. Arrows indicated extension of the 95^th^ percentile to the indicated values. n = 26-47 traces. * *P* < 0.05, ** *P* < 0.01. Two-tailed student’s *t*-test with Bonferroni’s correction compared to *h^WT^*. (**C**) Acetylcholine is the primary excitatory neurotransmitter in the CNS of *Drosophila.* Systemic injection of nicotinic acetylcholine receptor antagonist mecamylamine (MEC) blocks the spontaneous firing in *h^D33V^/h^R95H^* indicating the aberrant firing is generated from the CNS. (**D**) Firing pattern analyses. Plots of the instantaneous firing frequency (ISI^−1^) vs. the instantaneous coefficient of variation (CV_2_, a measure of firing regularity) readily distinguish grooming-related firing (*h^WT^* shown) from self-similar bursting in *h^D33V^* and *h^D33V^/h^R95H^*. Plots from representative firing patterns are shown (see Methods for details on construction of ISI^-1^-CV_2_ plots).

In contrast to the relatively mild phenotypes displayed by homozygous KI lines, we found clear, ongoing, seizure activity in *h^D33V^/h^R95H^* trans-heterozygotes, manifesting as high frequency spike burst discharges (Fig. 5A, Supplementary Video 2), with overall firing rates for *h^D33V^/h^R95H^* substantially greater than *h^WT^* or *h^WT^/h^R95H^* flies reared on the same media (Fig. 5B). To determine if this aberrant spiking activity originated from the CNS or endogenously from the motor unit, we blocked central excitatory neurotransmission and monitored the effect on spiking activity. In flies, acetylcholine is the primary excitatory neurotransmitter in the CNS [46], while glutamate is the transmitter at the neuromuscular junction [47]. We applied the nicotinic acetylcholine receptor blocker mecamylamine using a rapid systemic injection protocol [33], and found the spike discharges in *h^D33V^/h^R95H^* were abolished (Fig. 5C), indicating aberrant CNS activity drove the seizure-associated motor unit discharges in these flies.

During seizure-associated discharges in *h^D33V^/h^R95H^* mutants, bursts in *h^D33V^*, and grooming-associated activity in *h^WT^* and other KI flies, the DLM spiking intervals were highly variable, with instantaneous firing rates (defined as the reciprocal of inter-spike intervals, i.e. ISI^-1^) ranging from ~1 Hz to nearly 50 Hz. However, these activity patterns could be readily distinguished and were remarkably characteristic within mutant flies. To quantitatively delineate seizure-associated activity in KI flies from grooming related spiking, we employed a non-linear dynamical system approach we had previously used to describe firing patterns in the *sgll^95^* mutants and other hyperexcitable flies (Fig. 5D) [24,33]. For each spike in the recording, we plotted the spike’s instantaneous firing rate (ISI^−1^) against the instantaneous coefficient of variation (CV_2_, see Methods for definition). High CV_2_ values indicate irregular firing, while low values indicate rhythmic activity. Previously, we have shown seizure-related bursting in *sgll^95^* mutants corresponded to self-similar “looping” trajectory in the phase-space analysis, while grooming activity displayed a distinctive trajectory limited to high CV_2_ values (> 0.5). Based on the ISI^−1^-CV_2_ plots (Fig. 5D, representative trajectories shown), the orbiting trajectories of *h^D33V^* bursting, and *h^D33V^*/*h^R95H^* spike discharges were readily distinguished from grooming-related behavior and from one-another. Furthermore, the trajectories of *h^D33V^/h^R95H^* firing were qualitatively similar to spontaneous seizures in *sgll^95^* mutants [24], suggesting a shared neural mechanism underlying the aberrant activity.

### PLP is required for both development and function of adult brain

No *h^R95H^* homozygotes were generated on the normal diet (Fig. 3A), suggesting that the PNPO activity in these homozygous flies is insufficient to convert dietary VB6 to PLP to support development. We expected that PLP supplementation could rescue the developmental impairment. We therefore supplemented varying doses of PLP to breeders (Fig. 6A). Remarkably, we observed a partial rescue with 4 g/ml of PLP and a complete rescue with 40 or 400 *μ*g/ml of PLP (Fig. 6A).

**Figure 6.**
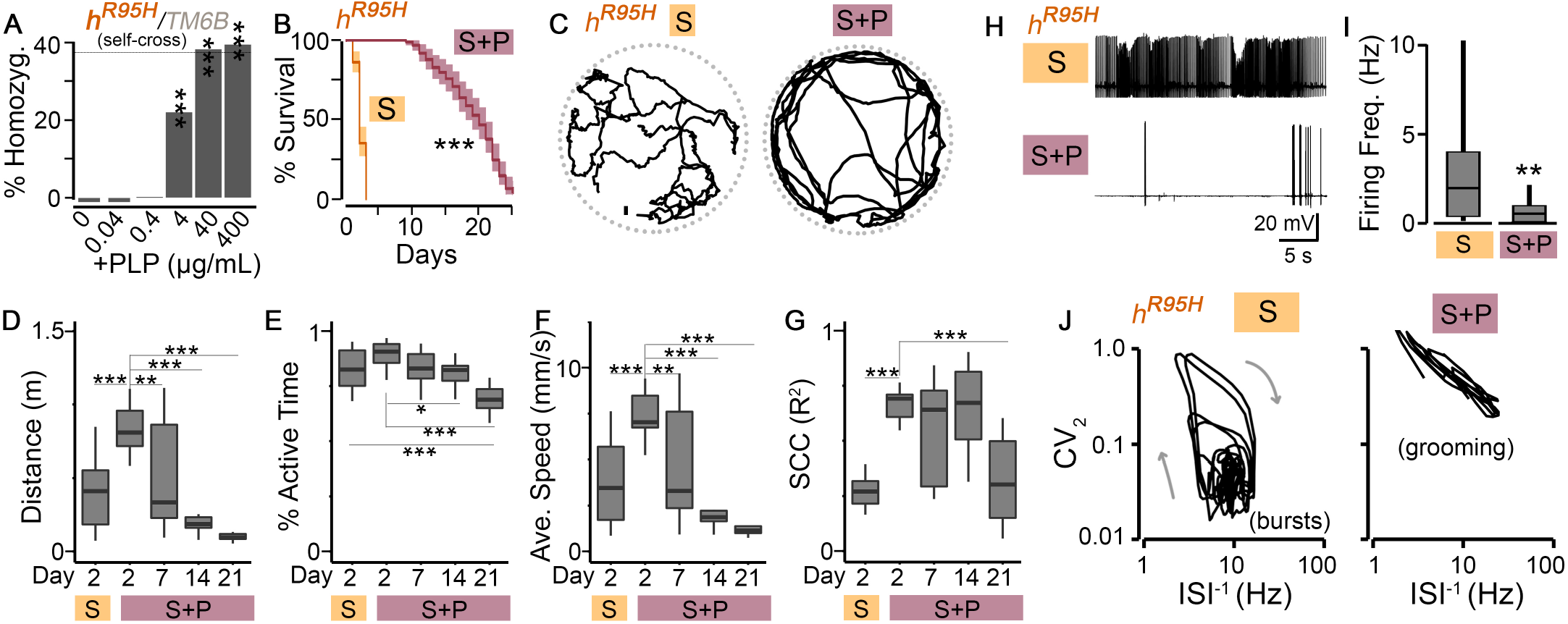
Development impairment, shortened lifespan, abnormal locomotion, and spontaneous seizure discharges in *h^R95H^* homozygotes are rescued by PLP supplementation. (**A**) Generation of *h^R95H^* homozygous flies from *h^R95H^/TM6B* with PLP supplementation. Chi-square test of homogeneity compared to the control group with 0 *μ*g/mL of PLP. n = 840-1163. The dotted gray line indicates the homozygosity ratio (% Homozygo.) from *h^WT^/TM6B* (Fig. **3A**) (**B**) Lifespan of *h^R95H^* homozygotes (generated with 400 *μ*g/mL of PLP) on sugar-only (S) or sugar supplemented with PLP (S+P, 400 *μ*g/ml). n = 100-110. Log-rank test. (**C**) Representative tracks during a 60-second interval of a single fly reared on S or S+P media for 2 days. (D-G) Quantification of *h^R95H^* walking over a 3-minute interval for flies reared on indicated media. n = 9-16 flies. * *P* < 0.05, ** *P* < 0.01, *** *P* < 0.001. One-way ANOVA with Tukey’s post hoc. (**D**) Total distance travelled. (**E**) Percentage of active time. (**F**) Average speed. (**G**) Speed correlation coefficient (SCC). (**H**) Representative traces of DLM firing from tethered *h^R95H^* flies. Note the burst discharges present in flies reared on sugar-only media are suppressed by PLP supplementation. (**I**) Average firing frequency. n = 24-33, ** *P* < 0.01, *** *P* < 0.001. Two-tailed student’s t-test. (**J**) Representative ISI^-1^-CV_2_ trajectories. Note that seizure-associated bursts are suppressed after PLP supplementation, and grooming (high CV_2_) associated patterns are observed.

To examine whether PLP is also indispensable for adult flies, we maintained *h^R95H^* homozygous flies (developed with PLP supplementation, 400 *μ* g/ml) on sugar with or without continued PLP supplementation (400 *μ*g/ml). With PLP supplementation *h^R95H^* mutants survived for several weeks (Fig. 6B, median lifespan: ~ 18 days). In contrast, without PLP supplementation, these flies could not survive for more than 3 days, demonstrating that PLP is also required for the survival of adult flies.

Similar to *h^D33V^*/*h^R95H^* (Fig. 5), *h^R95H^* homozygous flies also exhibited seizure-like behaviors before their death. To characterize these abnormal behaviors and to further study the role of PLP in the behavioral output, we monitored the locomotor activity of *h^R95H^* homozygotes with or without PLP supplementation (Fig. 6C, Supplementary Video 3). Compared to *h^R95H^* homozygotes without PLP supplementation, *h^R95H^* homozygotes with PLP supplementation for 2 days showed significantly improved levels in total distance traveled, percentage of active time, average speed, and SCC (Fig. 6D-G), demonstrating that PLP deficiency is responsible for the low behavioral performance of *h^R95H^* homozygous flies. The improvement of the behavioral performance declined with age, which correlates well with the increased lethality (Fig. 6B).

Consistent with our previous findings that PNPO deficiency leads to increased spontaneous firing and seizure discharges [24], *h^R95H^* homozygotes on sugar also exhibited clear spontaneous seizure discharges in the tethered fly preparation (Fig. 6H, Supplementary Video 4). Remarkably, these seizure-associated discharges were completely suppressed by PLP supplementation, as demonstrated by greatly reduced median firing rate (Fig. 6I, 1.9 Hz to 0.5 Hz for flies on sugar and sugar-only supplemented with PLP, respectively.) and the alteration from burst-associated trajectories to grooming-associated patterns in the ISI^-1^-CV_2_ plots (Fig. 6J).

Taken together, these results demonstrate that PLP is required for both development and normal brain function in adult flies.

### *h^WT^/h^R95H^* flies are susceptible to electroconvulsive seizures due to a DN effect of h^R95H^

We observed lethality from *h^WT^*/*h^R95H^* flies when reared on sugar (Fig. 3C). The lethality could be due to either haploinsufficiency or a DN effect of h^R95H^ on h^WT^. To test these two possibilities, we compared the survival rate of *h^WT^*/*h^R95H^* flies to that of *h^WT^*/*Df* flies in which the *sgll* gene is deleted in the *Df* line. Since there is only one *hPNPO* allele in *h^WT^/Df* flies, we reasoned that if the lethality was due to haploinsufficiency, survival of *h^WT^/h^R95H^* flies would be similar to *h^WT^/Df.* If the lethality was due to a DN effect, *h^WT^*/*h^R95H^* would exhibit worse survival than *h^WT^*/*Df* flies. As shown in Fig. 7A, the median survival rates are 19 and 27 days for *h^WT^/h^R95H^* and *h^WT^/Df*, respectively (P <0.001), demonstrating a DN effect of h^R95H^.

**Figure 7.**
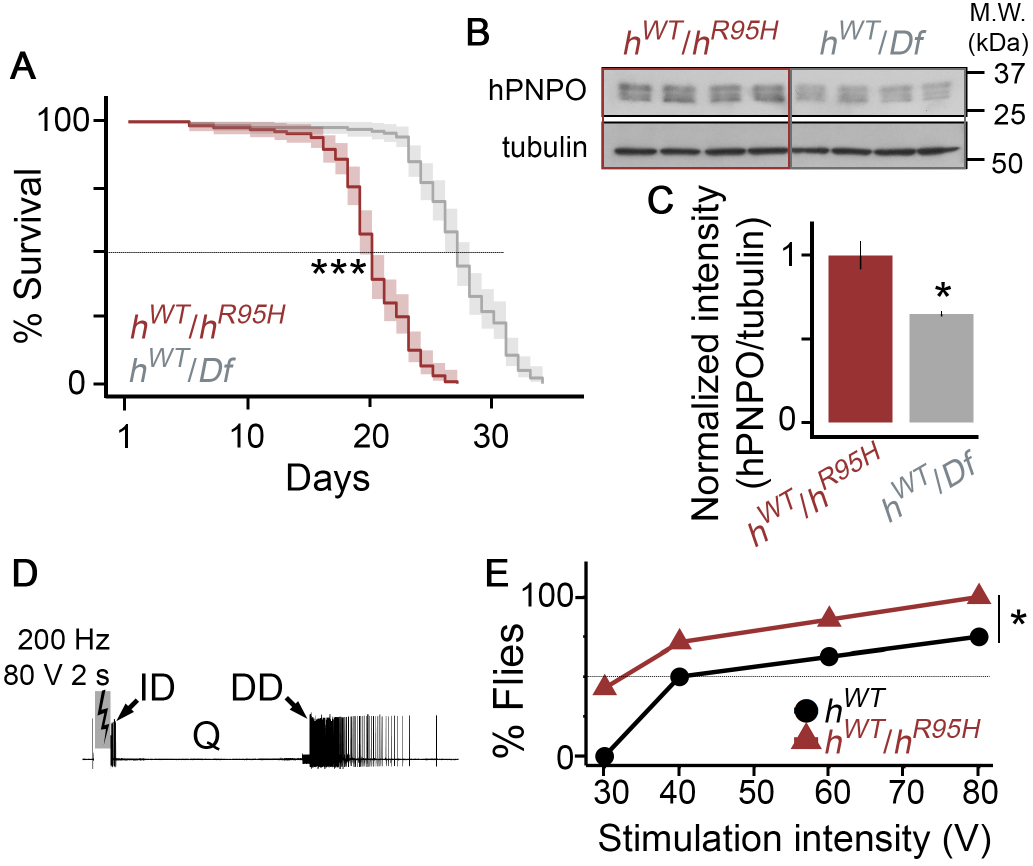
The *h^R95H^* allele has a dominant-negative effect on the *h^WT^* allele. (**A**) Survival of *h^WT^/h^R95H^* and *h^WT^/Df* flies on the sugar-only diet. n = 133-136. *** *P* < 0.001. Log-rank test. (**B**) Western blot of adult fly head homogenate from *h^WT^/h^R95H^* and *h^WT^/Df* flies. Tubulin is the loading control. n = 4 per genotype. (**C**) Quantifications of the intensity of *hPNPO* bands from four biological replicates in each genotype. All three bands were included in the quantification. Error bars represent Mean ±SEM. * *P* < 0.05. Two-tailed student’s t-test. (**D**) Electroconvulsive seizure (ECS) activity. High-frequency stimulation across the brain (200 Hz for 2 s) can trigger a stereotypic seizure activity sequence: an initial spike discharge (ID), a quiescent period (Q) and a delayed spike discharge (DD). (**E**) The fraction of individuals displaying DD as a function of stimulation intensity. n = 6-8. * *P* < 0.05, Kruskal-Wallis non-parametric ANOVA.

Functional PNPO is a dimerized protein [15], so the DN effect of h^R95H^ on h^WT^ was likely mediated through the formation of a heterodimer between them. However, the *hPNPO* protein level was decreased in *h^R95H^*/*sgll^+^* flies (Fig. 2A), suggesting that the monomer or homodimer of *h^R95H^* is unstable. How could the DN effect be possible if *h^R95H^* is not stable? One possibility was that h^R95H^ was stabilized by forming a heterodimer with h^WT^ in *h^WT^*/*h^R95H^* flies. The stabilization was not observed in *h^R95H^*/*sgll^+^* flies because *h^R95H^* is less likely to form a heterodimer with the *Drosophila PNPO* due to their differences in the length and amino acid sequences [10]. If h^R95H^ was indeed stabilized in *h^WT^/h^R95H^* flies, we would expect that the hPNPO protein level in *h^WT^*/*h^R95H^* to be twice as high as that in *h^WT^*/*Df* since the former has two *hPNPO* alleles and the latter only has one. That is exactly what we observed (Fig. 7B-C). Therefore, h^R95H^ exerts its DN effects by forming a heterodimer with h^WT^.

The DN effect would presumably make *h^WT^*/*h^R95H^* flies susceptible to seizures. However, we did not observe spontaneous seizures in them (Fig. 5A-B). We thus examined if they were prone to seizures upon high-frequency electrical stimulation (HFS). HFS across the brain can induce stereotypic electroconvulsive seizure (ECS) discharges in flies (Fig. 7D, Supplementary Video 5) [36,48]. Compared to control flies, seizure-prone flies usually require lower stimulation intensities to induce seizures [41,49]. Many seizure-prone mutants have thus been identified and characterized [36,41,43,44,48,50]. We found a substantial reduction in the ECS stimulation threshold in *h^WT^*/*h^R95H^* flies compared to *h^WT^* counterparts. When *h^WT^*/*h^R95H^* heterozygous and *h^WT^* homozygous flies were stimulated with a low voltage (30 V), seizures were already induced in 50% of *h^WT^*/*h^R95H^* flies while no seizures were observed in *h^WT^* (Fig. 7E). The increased sensitivity in *h^WT^/h^R95H^* was maintained when the stimulation intensity was gradually increased. At 80 V, essentially all *h^WT^/h^R95H^* flies showed ECS discharges while only about 60% of *h^WT^* were affected. Thus, the DN effect of *h^R95H^* confers the susceptibility of *h^WT^/h^R95H^* flies to seizures.

## Discussion

Neonatal epileptic encephalopathy patients with PNPO deficiency exhibit a wide spectrum of symptoms, including developmental impairments, intellectual disability, behavioral disorders, and seizures of various types, frequency and severity [3,18,20,51]. It has been suggested that both genetic and environmental factors contribute to the phenotypic variation [3]. Using a genetically tractable model system and well-controlled dietary conditions, we accomplished the functional and molecular characterization of human *PNPO* epilepsy-associated alleles in *Drosophila*. Our experimental evidence indicates that *PNPO* mutations, diet, and allele-diet interactions all contribute to the final phenotype expression.

Our results demonstrate that the diet’s impact on the phenotype expression depends on *PNPO* alleles, and it is more impactful when PNPO deficiency is mild. For example, with different dietary breeding and testing conditions, *h^R116Q^*, *h^D33V^*, or *h^WT^/h^R95H^* flies exhibit different behavioral outcomes, whereas *h^D33V^/h^R95H^* have relatively invariant phenotypes (Fig. 4, Supplementary Fig. 5). A less variable phenotype was also observed in patients carrying severe PNPO mutations. The seizure onsets in patients with severe PNPO mutations is ranged from hours to weeks compared to hours to years in patients with mild mutations [17]. The less significant effect of diet in KI flies or patients with severe PNPO deficiency is likely due to the diminished capability of mutant PNPOs in converting dietary VB6 to PLP. On the other hand, although mild PNPO deficiency has the capacity to convert dietary VB6 to PLP, and it is less affected by dietary conditions to cause acute effects, the fact that *h^R116Q^* and *h^D33V^*flies exhibit shortened lifespans on a normal diet (Fig. 3B) suggests that the deleterious effect of mild PNPO deficiency can be cumulative. R116Q is found in ~ 10% of the general population, and 1% of the general population are homozygous for R116Q [52]. Whether mild PNPO deficiency caused by R116Q may be exacerbated by other genetic and/or environmental factors to cause epilepsy and/or other diseases in a long-time period remains to be investigated.

Developmental delay is one common symptom manifested by PNPO deficient patients [3,18,20,53], raising the question of whether specific developmental impairments lead to seizures. By providing flies with PLP during either the developmental or adult stage, we demonstrate that developmental impairments and seizures in adults are dissociable (Fig. 6). Given PNPO is ubiquitously expressed [12], it remains to be determined whether the lethality foci involve solely the brain or additional organs and tissues. Further functional studying of *PNPO* can bring insights into the mechanistic links to other comorbidities (e.g., intellectual disability) manifested in PNPO deficient patients.

We show with *in vivo* evidence that R116Q is a mild mutation. The molecular and functional consequences of R116Q have been controversial. R116Q was first reported in patients with neonatal epileptic encephalopathy [3], suggesting it is likely a severe mutation. Later clinical reports show that unlike seizures in patients carrying other PNPO mutations, seizures in R116Q patients can occur beyond the neonatal stage [3,4,5], indicating that R116Q is more likely to be a relatively mild mutation. The molecular consequence of R116Q is also inconclusive. An earlier *in vitro* study shows that R116Q decreases the enzyme activity to 83% of the WT protein [3]. In contrast, two recent studies report decreases to ~ 40% and ~ 1% of the WT protein instead [4,41]. We found that *h^R116Q^* flies develop well (Fig. 3A) and do not show spontaneous seizures (Fig. 5A-B), consistent with a mild defect of the enzyme function. Nevertheless, the finding that R116Q alters PNPO protein localization in the brain (Fig. 2D) awaits further investigation into how this specific molecular change contributes to the various phenotypes seen in *h^R116Q^* (Fig. 4, Supplementary Fig. 5).

We find unique DLM burst firings in *h^D33V^* flies. Coincidentally, distinct EEG patterns have been reported in patients carrying the D33V mutation, who often exhibit temporal sharp waves or multifocal sharp waves instead of burst suppression in the majority of PNPO deficient patients [51]. It is unclear whether the unique firing pattern in *h^D33V^* flies share common mechanisms with the distinct EEG patterns in D33V patients. The amino acid D33 (Aspartate 33) is located in the N-terminus of the PNPO protein [12]. The N-terminus of PNPO is often not retained in crystal samples [15], posing challenges in examining D33V in structural studies. Our molecular studies in *h^D33V^* KI flies demonstrate a decrease in the mRNA level and, consequently, the protein level of PNPO (Fig. 2A-C). However, it seems unlikely that the decreased PNPO protein level is responsible for the distinct firing in *h^D33V^* flies since a decreased PNPO protein level is also observed in *h^R95H^* flies which exhibit regular seizure discharges (Fig. 6H-J). Future studies should investigate whether D33V mutant protein alters neuronal functions in unique ways.

Neonatal epileptic encephalopathy caused by PNPO deficiency is thought to be an autosomal recessive disease (OMIM #610090), implying that PNPO mutant carriers do not show any overt phenotype. Indeed, in all reported PNPO deficiency cases, 78% are homozygous for a specific mutation, and 21% are compound heterozygotes [17], except for one patient reported to be a heterozygote [3]. Yet, in KI flies, we found a readily detectable DN effect from the *h^R95H^* allele (Fig. 7). The DN effect is potentially associated with heterodimer formation between h^R95H^ and h^WT^. Structural studies have shown that amino acid R95 (Arginine 95) is required for the co-factor FMN binding [15,54] and thus PNPO enzyme activity [14,15]. An altered FMN binding site may lead to global conformation changes and hence a partially or fully inactive enzyme. It is tempting to predict that mutations that do not affect dimerization but affect FMN binding may have a DN effect. Therefore, human carriers of certain PNPO mutations could be susceptible to PNPO deficiency-induced epilepsy and other diseases.

In summary, our data has important implications in understanding the genetics of PNPO deficiency and its clinical implications. Our studies demonstrate the role of *PNPO*allele-diet interaction in the phenotype expression, uncover a diversity of molecular effects of *PNPO* variants, and highlight the contribution of PNPO deficiency to epilepsy in general. These studies also demonstrate that KI *Drosophila* models are valuable for systematically analyzing the functional and molecular effects of each *PNPO* allele identified in epilepsy patients.

## Acknowledgments

We thank Vytas Bindokas and Zijie Zhang for technical support. We thank Benjamin Wang for discussions. We thank the Bloomington Drosophila Stock Center for *Drosophila* stocks and Fungene Drosophila Resource Center for generating KI flies.

## Supplementary material

Supplementary material is available online.

## References

[1] Marcelina Parra, Seth Stahl, and Hanjo Hellmann. Vitamin b6 and its role in cell metabolism and physiology. Cells, 7:84–112, 7 2018.

[2] Philippa B. Mills, Robert A H Surtees, Michael P. Champion, Clare E. Beesley, Neil Dalton, Peter J. Scamber, Simon J R Heales, Anthony Briddon, Irene Scheimberg, Georg F. Hoffmann, Johannes Zschocke, and Peter T. Clayton. Neonatal epileptic encephalopathy caused by mutations in the pnpo gene encoding pyridox(am)ine 5-phosphate oxidase. Human Molecular Genetics, 14:1077–1086, 2005.

[3] Philippa B. Mills, Stephane S M Camuzeaux, Emma J. Footitt, Kevin A. Mills, Paul Gissen, Laura Fisher, Krishna B. Das, Sophia M. Varadkar, Sameer Zuberi, Robert McWilliam, T. Stodberg, Barbara Plecko, Matthias R. Baumgartner, Oliver Maier, Sophie Calvert, Kate Riney, Nicole I. Wolf, John H. Livingston, Pronab Bala, Chantal F. Morel, Franôois Feillet, Francesco Raimondi, En-nio Del Giudice, W. Kling Chong, Matthew Pitt, and Peter T. Clayton. Epilepsy due to pnpo mutations: genotype, environment and treatment affect presentation and outcome. Brain, 137:1350–1360, 5 2014.

[4] Martino L. di Salvo, Mario Mastrangelo, Isabel Nogués, Manuela Tolve, Alessandro Paiardini, Carla Carducci, Davide Mei, Martino Montomoli, Angela Tramonti, Renzo Guerrini, Roberto Contestabile, and Vincenzo Leuzzi. Pyridoxine-5-phosphate oxidase (pnpo) deficiency: Clinical and biochemical alterations associated with the c.347g >a (p: Arg116gln) mutation. Molecular Genetics and Metabolism, 122:135–142, 9 2017.

[5] Jiao Xue, Xingzhi Chang, Yuehua Zhang, and Zhixian Yang. Novel phenotypes of pyridox(am)ine-5’-phosphate oxidase deficiency and high prevalence of c.445_448del mutation in chinese patients. Metabolic Brain Disease, 32:1081–1087, 2017.

[6] Jacques L. Michaud, Mathieu Lachance, Fadi F. Hamdan, Lionel Carmant, Anne Lortie, Paola Diadori, Philippe Major, Inge A. Meijer, Emmanuelle Lemyre, Patrick Cossette, Heather C. Mefford, Guy A. Rouleau, and Elsa Rossignol. The genetic landscape of infantile spasms. Human Molecular Genetics, 23:4846–4858, 2014.

[7] The Interntional League Against Epilpesy Consortium on Complex Epilepsies. Genome-wide mega-analysis identifies 16 loci and highlights diverse biological mechanisms in the common epilepsies. Nature Communications, 9:5269, 12 2018.

[8] Riccardo Percudani and Alessio Peracchi. A genomic overview of pyridoxal-phosphate-dependent enzymes. supplementary material. EMBO reports, 4:850–4, 2003.

[9] Martin K. Safo, Irimpan Mathews, Faik N. Musayev, Martino L. Di Salvo, Daniel J. Thiel, Donald J. Abraham, and Verne Schirch. X-ray structure of escherichia coli pyridoxine 5’-phosphate oxidase complexed with fmn at 1.8 Å resolution. Structure, 8:751–762, 2000.

[10] Wanhao Chi, Li Zhang, Wei Du, and Xiaoxi Zhuang. A nutritional conditional lethal mutant due to pyridoxine 5’-phosphate oxidase deficiency in drosophila melanogaster. G3; Genes,Genomes,Genetics, 4:1147–1154, 2014.

[11] Po Yuan Chen, Hung Chi Tu, Verne Schirch, Martin K. Safo, and Tzu Fun Fu. Pyridoxamine supplementation effectively reverses the abnormal phenotypes of zebrafish larvae with pnpo deficiency. Frontiers in Pharmacology, 10:1–13, 2019.

[12] Jeong Han Kang, Mi Lim Hong, Dae Won Kim, Jinseu Park, Tae Cheon Kang, Moo Ho Won, Nam In Baek, Byung Jo Moon, Soo Young Choi, and Oh Shin Kwon. Genomic organization, tissue distribution and deletion mutation of human pyridoxine 5’-phosphate oxidase. European Journal of Biochemistry, 271:2452–2461, 2004.

[13] Martino Di Salvo, Emily Yang, Genshi Zhao, Malcolm E. Winkler, and Verne Schirch. Expression, purification, and characterization of recombinant escherichia coil pyridoxine 5’-phosphate oxidase. Protein Expression and Purification, 1998.

[14] Jorge E. Churchich. Brain pyridoxine-5’-phosphate oxidase: A dimeric enzyme containing one fmn site. European Journal of Biochemistry, 138:327–332, 1984.

[15] Faik N Musayev, Martino L Di Salvo, Tzu-Ping Ko, Verne Schirch, and Martin K Safo. Structure and properties of recombinant human pyridoxine 5’-phosphate oxidase. Protein science, 12:1455–1463, 2003.

[16] Jung-Do Choi, Delores M. Bowers-Komro, Michael D. Davis, Dale E. Edmondson, and Donald B. McCormick. Kinetic properties of pyridoxamine (pyridoxine)-5’-phosphate oxidase from rabbit liver. Journal of Biological Chemistry, 258:840–845, 1983.

[17] Wanhao Chi. Genetic and neural mechanisms of pnpo deficiency in vitamin b6-dependent epilepsy. PhD Thesis, The University of Chicago, 2019.

[18] Malak Alghamdi, Fahad A. Bashiri, Marwa Abdelhakim, Nouran Adly, Dima Z. Jamjoom, Khalid M. Sumaily, Bandar Alghanem, and Stefan T. Arold. Phenotypic and molecular spectrum of pyridoxamine-5-phosphate oxidase deficiency: A scoping review of 87 cases of pyridoxamine-5-phosphate oxidase deficiency. Clinical Genetics, 2021.

[19] Morad Khayat, Stanley H. Korman, Pnina Frankel, Zalman Weintraub, Sylvia Hershckowitz, Vered Fleisher Sheffer, Mordechai Ben Elisha, Ronald A. Wevers, and Tzipora C. Falik-Zaccai. Pnpo deficiency: An under diagnosed inborn error of pyridoxine metabolism. Molecular Genetics and Metabolism, 94:431–434, 2008.

[20] Alina Levtova, Stephane Camuzeaux, Anne-Marie Laberge, Pierre Allard, Catherine Brunel-Guitton, Paola Diadori, Elsa Rossignol, Keith Hyland, Peter T Clayton, Philippa B Mills, and Grant A Mitchell. Normal cerebrospinal fluid pyridoxal 5’-phosphate level in a pnpo-deficient patient with neonatal-onset epileptic encephalopathy. JIMD reports, 22:67–75, 2015.

[21] Mohini S. Ghatge, Mohammed Al Mughram, Abdelsattar M. Omar, and Martin K. Safo. Inborn errors in the vitamin b6 salvage enzymes associated with neonatal epileptic encephalopathy and other pathologies. Biochimie, 183:18–29, 2021.

[22] Anna Barile, Isabel Nogués, Martino L. di Salvo, Victoria Bunik, Roberto Contestabile, and Angela Tramonti. Molecular characterization of pyridoxine 5-phosphate oxidase and its pathogenic forms associated with neonatal epileptic encephalopathy. Scientific Reports, 10:1–15, 2020.

[23] Xu Zhang, Wouter H. Koolhaas, and Frank Schnorrer. A versatile two-step crispr- and rmce-based strategy for efficient genome engineering in drosophila. G3: Genes, Genomes, Genetics, 2014.

[24] Wanhao Chi, Atulya S R Iyengar, Monique Albersen, Marjolein Bosma, Nanda M Verhoeven-Duif, Chun-Fang Wu, and Xiaoxi Zhuang. Pyridox (am) ine 5’-phosphate oxidase deficiency induces seizures in drosophila melanogaster. Human Molecular Genetics, 28:3126–3136, 9 2019.

[25] Scott J. Gratz, Fiona P. Ukken, C. Dustin Rubinstein, Gene Thiede, Laura K. Donohue, Alexander M. Cummings, and Kate M. Oconnor-Giles. Highly specific and efficient crispr/cas9-catalyzed homology-directed repair in drosophila. Genetics, 196:961–971, 2014.

[26] Andrew R. Bassett, Charlotte Tibbit, Chris P. Ponting, and Ji-Long Liu. Highly efficient targeted mutagenesis of drosophila with the crispr/cas9 system. Cell reports, 4:220–8, 7 2013.

[27] A.W.K. Frankel and Jr. G.E. Brosseau. A drosophila medium that does not require dried yeast. Drosophila. Inf. Service, 43:184, 1968.

[28] Barret D Pfeiffer, Arnim Jenett, Ann S Hammonds, Teri-T B Ngo, Sima Misra, Christine Murphy, Audra Scully, Joseph W Carlson, Kenneth H Wan, Todd R Laverty, et al. Tools for neuroanatomy and neurogenetics in drosophila. Proceedings of the National Academy of Sciences, 105(28):9715–9720, 2008.

[29] Johannes Schindelin, Ignacio Arganda-Carreras, Erwin Frise, Verena Kaynig, Mark Longair, Tobias Pietzsch, Stephan Preibisch, Curtis Rueden, Stephan Saalfeld, Benjamin Schmid, Jean Yves Tinevez, Daniel James White, Volker Hartenstein, Kevin Eliceiri, Pavel Tomancak, and Albert Cardona. Fiji: An open-source platform for biological-image analysis. Nature Methods, 9:676–682, 2012.

[30] Atulya Iyengar, Jordan Imoehl, Atsushi Ueda, Jeffery Nirschl, and Chun Fang Wu. Automated quantification of locomotion, social interaction, and mate preference in drosophila mutants. Journal of Neurogenetics, 26:306–316, 2012.

[31] Atulya Iyengar and Chun Fang Wu. Flight and seizure motor patterns in drosophila mutants: Simultaneous acoustic and electrophysiological recordings of wing beats and flight muscle activity. Journal of Neurogenetics, 28:316–328, 2014.

[32] G. R. Holt, W. R. Softky, C. Koch, and R. J. Douglas. Comparison of discharge variability in vitro and in vivo in cat visual cortex neurons. Journal of Neurophysiology, 75:1806–1814, 5 1996.

[33] Jisue Lee, Atulya Iyengar, and Chun-Fang Wu. Distinctions among electroconvulsion- and proconvulsant-induced seizure discharges and native motor patterns during flight and grooming: quantitative spike pattern analysis in drosophila flight muscles. Journal of Neurogenetics, 33:125–142, 4 2019.

[34] Iris C. Howlett and Mark A. Tanouye. Seizure-sensitivity in drosophila is ameliorated by dorsal vessel injection of the antiepileptic drug valproate. Journal of Neurogenetics, 27:143–150, 2013.

[35] Jing W. Wang, Allan M. Wong, Jorge Flores, Leslie B. Vosshall, and Richard Axel. Two-photon calcium imaging reveals an odor-evoked map of activity in the fly brain. Cell, 112:271–282, 2003.

[36] Jisue Lee and Chun-Fang Wu. Electroconvulsive seizure behavior in drosophila: analysis of the physiological repertoire underlying a stereotyped action pattern in bang-sensitive mutants. The Journal of Neuroscience, 22:11065–11079, 2002.

[37] Susan E. Celniker, Laura A.L. L Dillon, Mark B. Gerstein, Kristin C. Gunsalus, Steven Henikoff, Gary H. Karpen, Manolis Kellis, Eric C. Lai, Jason D. Lieb, David M. Macalpine, Gos Micklem, Fabio Piano, Michael Snyder, Lincoln Stein, Kevin P. White, and Robert H. Waterston. Unlocking the secrets of the genome. Nature, 459:927–930, 2009.

[38] Aoife Larkin, Steven J. Marygold, Giulia Antonazzo, Helen Attrill, Gilberto dos Santos, Phani V. Garapati, Joshua L. Goodman, L. Sian Gramates, Gillian Millburn, Victor B. Strelets, Christopher J. Tabone, and Jim Thurmond. Flybase: Updates to the drosophila melanogaster knowledge base. Nucleic Acids Research, 49:D899–D907, 1 2021.

[39] Dhananjay A. Wagh, Tobias M. Rasse, Esther Asan, Alois Hofbauer, Isabell Schwenkert, Heike Dürrbeck, Sigrid Buchner, Marie Christine Dabauvalle, Manuela Schmidt, Gang Qin, Carolin Wichmann, Robert Kittel, Stephan J. Sigrist, and Erich Buchner. Bruchpilot, a protein with homology to elks/cast, is required for structural integrity and function of synaptic active zones in drosophila. Neuron, 49:833–844, 2006.

[40] Kei Ito, Kazunori Shinomiya, Masayoshi Ito, J. Douglas Armstrong, George Boyan, Volker Hartenstein, Steffen Harzsch, Martin Heisenberg, Uwe Homberg, Arnim Jenett, Haig Keshishian, Linda L. Restifo, Wolfgang Rössler, Julie H. Simpson, Nicholas J. Strausfeld, Roland Strauss, and Leslie B. Vosshall. A systematic nomenclature for the insect brain. Neuron, 81:755–765, 2 2014.

[41] Daniel Kuebler and Mark A Tanouye. Modifications of seizure susceptibility in drosophila. Journal of neurophysiology, 83:998–1009, 2 2000.

[42] Ping Wang, Sudipta Saraswati, Zhuo Guan, Carol J. Watkins, Richard J. Wurtman, and J. Troy Littleton. A drosophila temperature-sensitive seizure mutant in phosphoglycerate kinase disrupts atp generation and alters synaptic function. Journal of Neuroscience, 24:4518–4529, 5 2004.

[43] Jisue Lee and Chun-Fang. Wu. Genetic modifications of seizure susceptibility and expression by altered excitability in drosophila na+ and k+ channel mutants. Journal of Neurophysiology, 96:2465–2478, 2006.

[44] Tim Fergestad, Bret Bostwick, and Barry Ganetzky. Metabolic disruption in drosophila bang-sensitive seizure mutants. Genetics, 173:1357–1364, 2006.

[45] Garrett A Kaas, Junko Kasuya, Patrick Lansdon, Atsushi Ueda, Atulya Iyengar, Chun-Fang Wu, and Toshihiro Kitamoto. Lithium-responsive seizure-like hyperexcitability is caused by a mutation in the drosophila voltage-gated sodium channel gene paralytic. eNeuro, 3:221–16, 11 2016.

[46] Paul M. Salvaterra and Toshihiro Kitamoto. Drosophila cholinergic neurons and processes visualized with gal4/uas-gfp. Gene Expression Patterns, 1:73–82, 2001.

[47] L. Y. Jan and Y. N. Jan. L-glutamate as an excitatory transmitter at the drosophila larval neuromuscular junction. The Journal of Physiology, 262:215–236, 1976.

[48] P Pavlidis and M A Tanouye. Seizures and failures in the giant fiber pathway of drosophila bang-sensitive paralytic mutants. The Journal of neuroscience, 15:5810–9, 1995.

[49] Louise Parker, Iris C. Howlett, Zeid M. Rusan, and Mark A. Tanouye. Seizure and epilepsy: Studies of seizure disorders in drosophila. International Review of Neurobiology, pages 1–21, 2011.

[50] Richard Marley and Richard a Baines. Increased persistent na+ current contributes to seizure in the slamdance bang-sensitive drosophila mutant. J. Neurophysiol., 106:18–29, 2011.

[51] Andrea Guerin, Aly. S. Aziz, Carly. Mutch, J.illian Lewis, Cristina. Y. Go, and Saadet. Mercimek-Mahmutoglu. Pyridox(am)ine-5-phosphate oxidase deficiency treatable cause of neonatal epileptic encephalopathy with burst suppression: Case report and review of the literature. Journal of Child Neurology, 30:1218–1225, 2014.

[52] Konrad J. Karczewski, Laurent C. Francioli, Grace Tiao, Beryl B. Cummings, Jessica Alföldi, Qingbo Wang, Ryan L. Collins, Kristen M. Laricchia, Andrea Ganna, Daniel P. Birnbaum, Laura D. Gauthier, Harrison Brand, Matthew Solomonson, Nicholas A. Watts, Daniel Rhodes, Moriel Singer-Berk, Eleina M. England, Eleanor G. Seaby, Jack A. Kosmicki, Raymond K. Walters, Katherine Tashman, Yossi Farjoun, Eric Banks, Timothy Poterba, Arcturus Wang, Cotton Seed, Nicola Whiffin, Jessica X. Chong, Kaitlin E. Samocha, Emma Pierce-Hoffman, Zachary Zappala, Anne H. O’Donnell-Luria, Eric Vallabh Minikel, Ben Weisburd, Monkol Lek, James S. Ware, Christopher Vittal, Irina M. Armean, Louis Bergelson, Kristian Cibulskis, Kristen M. Connolly, Miguel Covarrubias, Stacey Donnelly, Steven Ferriera, Stacey Gabriel, Jeff Gentry, Namrata Gupta, Thibault Jeandet, Diane Kaplan, Christopher Llanwarne, Ruchi Munshi, Sam Novod, Nikelle Petrillo, David Roazen, Valentin Ruano-Rubio, Andrea Saltzman, Molly Schleicher, Jose Soto, Kathleen Tibbetts, Charlotte Tolonen, Gordon Wade, Michael E. Talkowski, Benjamin M. Neale, Mark J. Daly, and Daniel G. MacArthur. The mutational constraint spectrum quantified from variation in 141,456 humans. Nature, 581:434–443, 5 2020.

[53] Licia Lugli, Maria Carolina Bariola, Luca Ori, Laura Lucaccioni, Alberto Berardi, and Fabrizio Ferrari. Further delineation of pyridoxine-responsive pyridoxine phosphate oxidase deficiency epilepsy: Report of a new case and review of the literature with genotype-phenotype correlation. Journal of Child Neurology, 34:937–943, 12 2019.

[54] Mohini S. Ghatge, Sayali S. Karve, Tanya M S David, Mostafa H. Ahmed, Faik N. Musayev, Kendra Cunningham, Verne Schirch, and Martin K. Safo. Inactive mutants of human pyridoxine 5-phosphate oxidase: a possible role for a noncatalytic pyridoxal 5-phosphate tight binding site. FEBS Open Bio, 6:398–408, 5 2016.

