## Supplementa figures & tables for "Diet modifies allele-specific phenotypes in *Drosophila* carrying epilepsy-associated *PNPO* variants"

### Methods

#### *Drosophila* husbandry

Behavioral recordings in Supplementary Fig. 5 F-J were performed on flies bred on Frankel & Brosseau's (FB) media [1] at the University of Iowa. For all other experiments, flies were generated on standard Cornmeal-Yeast-Molasses (CYM) media from the Fly Kitchen at the University of Chicago. Flies used in all experiments were raised and tested at room temperature ($\sim$ 23 $^{o}$C) in a 12:12 hour light:dark cycle. The deficiency lines *Df(3R)BSC221/TM6B* and *Df(3R)ED5223/TM6C* were obtained from the Bloomington Drosophila Stock Center (BDSC #9698, #9076).

#### Complementation test

Female flies, either homozygous ($h^{WT}$, $h^{R116Q}$, and $h^{D33V}$) or heterozygous ($h^{R95H}$/*TM6B*), and male flies from one of the *Df* strains were picked for breeding. A cohort of two to three such flies per sex was set up for each combination. F1 flies eclosed within six days from each cross were examined for the Balancer marker.

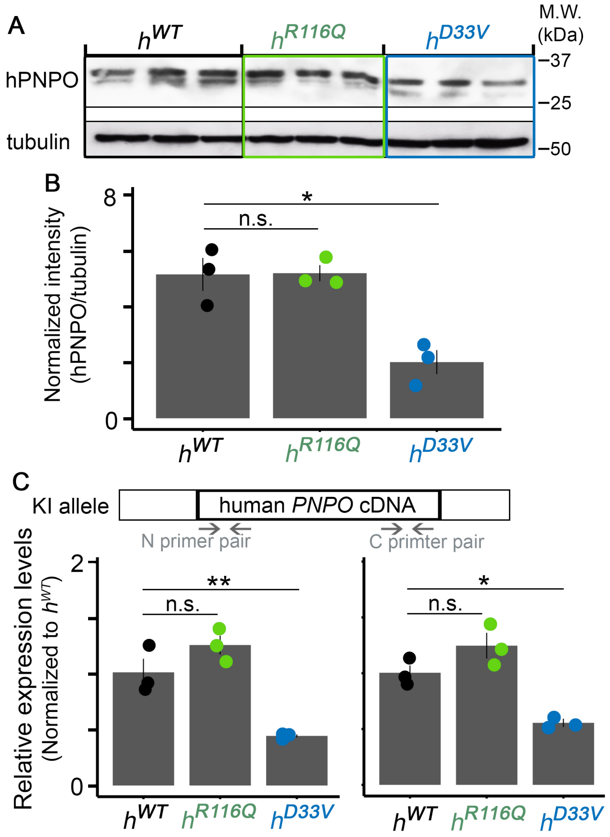


**Supplementary Figure 1. The *hPNPO* mRNA and protein levels in KI homozygotes.** (**A**) Western blot of adult fly head homogenate from $h^{WT}$, $h^{R116Q}$, and $h^{D33V}$ homozygotes. n = 3 per genotype. Tubulin is the loading control. (**B**) Quantifications of hPNPO protein level in panel A (all bands). (C) Expression of hPNPO at the mRNA level. n = 3 per genotype. Error bars represent Mean ± SEM. n.s. $P > 0.05$, * $P < 0.05$, ** $P < 0.01$. Two-tailed student's *t*-test with Bonferroni's correction.

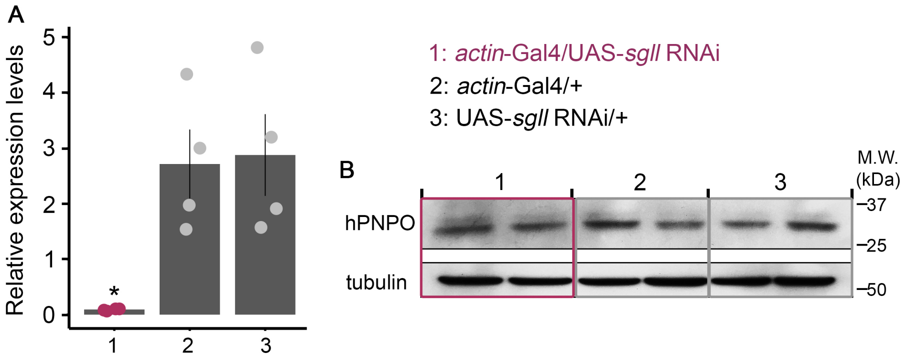


**Supplementary Figure 2. The antibody used for detecting hPNPO does not seem to recognize *Drosophila* PNPO.** (**A**) The *sgll* mRNA levels in ubiquitous *sgll* knockdown (genotype: *actin*-Gal4/UAS-*sgll* RNAi) and control flies (genotypes: *actin*-Gal4/+ and UAS-*sgll* RNAi/+). n = 4 per genotype. Error bars represent Mean ± SEM. * $P < 0.05$. One-way ANOVA with Tukey's post hoc. (**B**) Western blots from adult head homogenates with various genotypes. n = 2 per genotype. Tubulin was the loading control. One bind was detected from all three genotypes. The size of this band seems to be correct; the predicted molecular weight for *Drosophila* PNPO (~ 27 KDa). However, the band intensity in *sgll* knockdown flies is as same as that in two controls, indicating that this band is less likely to be *Drosophila* PNPO.

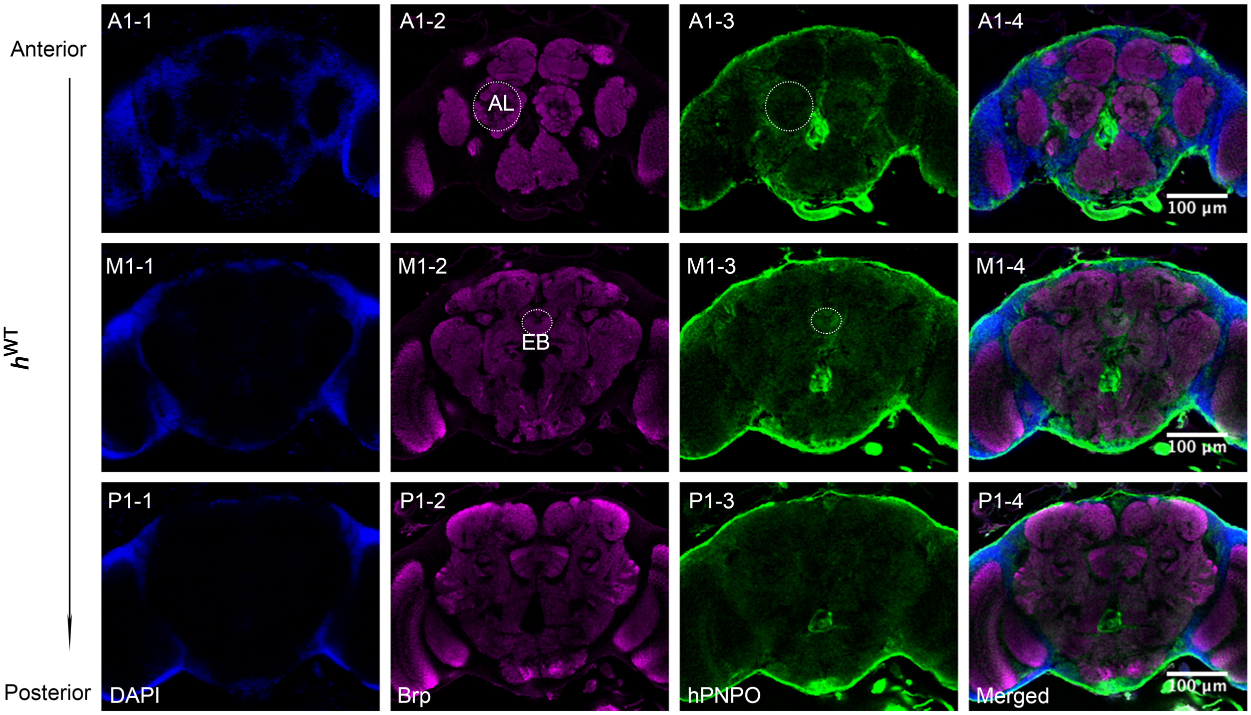


**Supplementary Figure 3. Expression pattern of $h^{WT}$ in the adult brain.** $h^{WT}$ was ubiquitously expressed in the brain with the strongest staining from cell body rind, a structure equivalent to the cortex in mammals [2]. Relatively strong staining was also observed in the areas surrounding neuropils, e.g., antennal lobe (AL) and ellipsoid body (EB). There was little overlap between Brp and hPNPO staining, suggesting that $h^{WT}$ is not enriched in the terminal structure. Blue, DAPI; Magenta, Brp; Green, hPNPO.

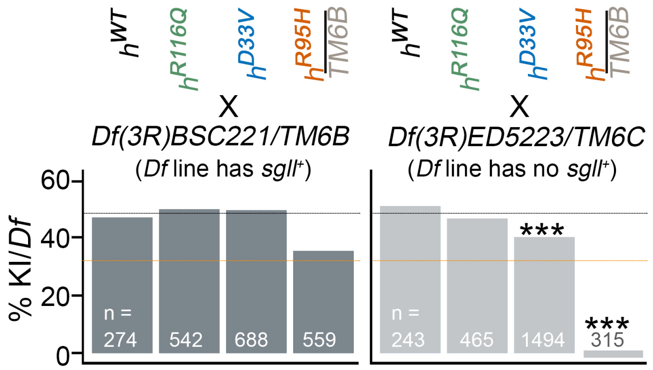


**Supplementary Figure 4. Complementation test.** Two Deficiency (*Df*) lines were used: one experimental line (genotype: *Df(3R)ED5223TM6C,Sb*) with the *sgll* gene deleted and one control line (genotype: *Df(3R)BSC221TM6B,Tb*) with a chromosome deletion adjacent to that of the experimental line. Each *Df* line was individually crossed with either homozygous flies from $h^{WT}$, $h^{R116Q}$, and $h^{D33V}$ lines or heterozygous flies from $h^{R95H}$ (due to homozygous lethality). Two dotted lines represent the expected ratio of KI/*Df* from breeding with three homozygous lines and the $h^{R95H}$/*TM6B* line, respectively. *** $P < 0.001$, Chi-square test of homogeneity compared to the expected value.

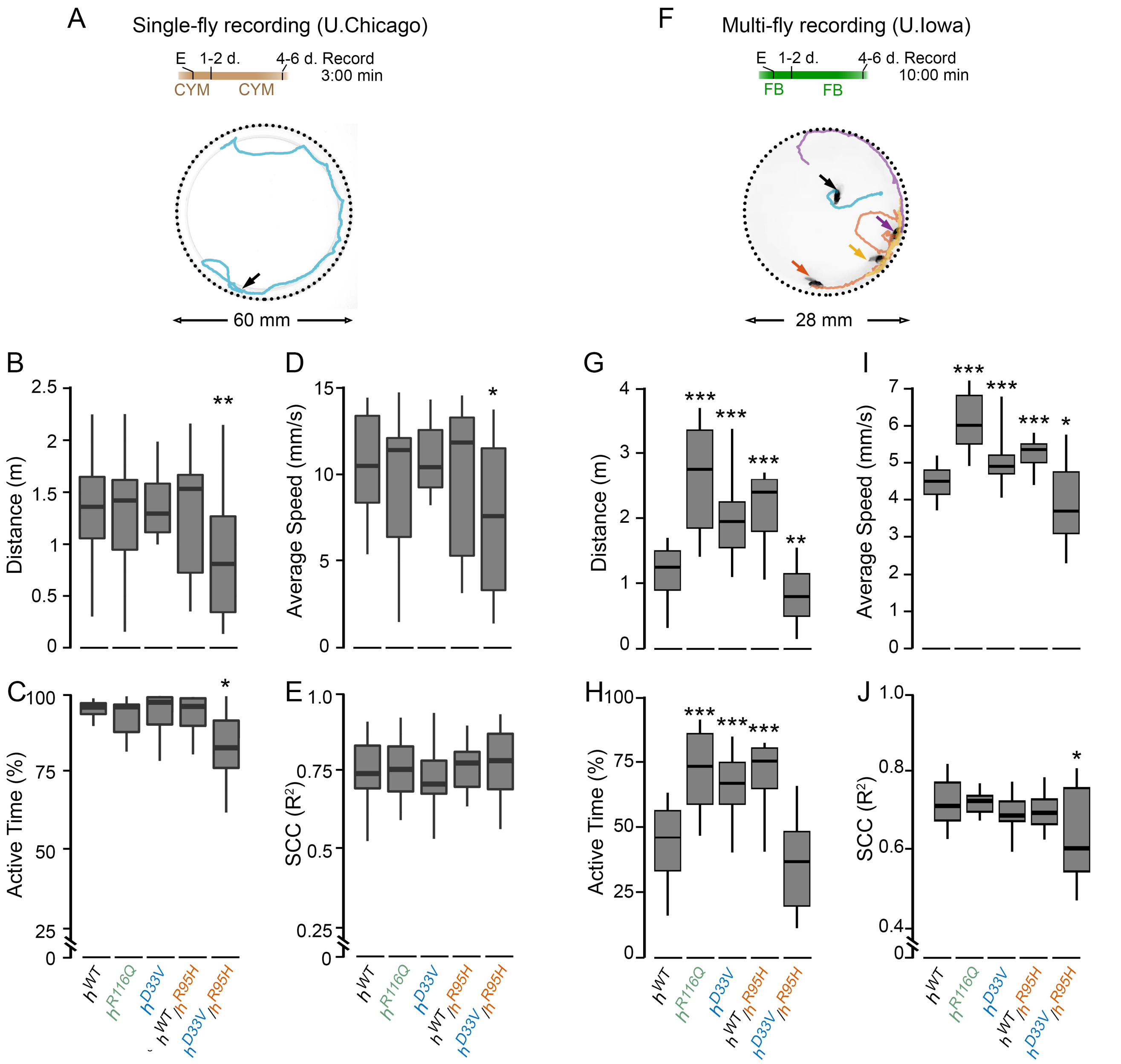


**Supplementary Figure 5. Behavioral analyses of KI flies on the standard diets.** (**A, F**) Breeding and testing conditions. Flies were eclosed (E) and maintained on either CYM or FB diet. (**B-E**) Total travelled distance, average speed, percentage of active time, and speed correlation coefficient (SCC) from flies in condition A. n = 18-24. (**G-J**) Total travelled distance, average speed, percentage of active time, and SCC from flies in condition F. n = 32-40. * $P < 0.05$, ** $P < 0.01$, ** $P < 0.001$. Two-tailed student's *t*-test with Bonferroni's correction compared to $h^{WT}$.

**Supplementary Table 1. gRNA and PCR primer sequences**

| | | |
|---|---|---|
| Generation of KI flies | gRNA-1-1 | GTGGCACCTCCGAGGCACCTTGG |
| | gRNA-1-2 | GTTCCACAGAGTTACCAGCAGG |
| | gRNA-2-1 | GAACCAGTTTGCTGTCGACTTCGG |
| | gRNA-2-2 | GGTTGGGTGTACGAACGGCTGG |
| KI mutation confirmation | Primer Forward | TGAAGTTGCTGCAAACAATTCGAAGG |
| | Primer Reverse | CTAAGGTGCAAGTCTCTCATAGAGCC |
| qPCR primers | KI-N Forward | ATGACGTGCTGGCTGCG |
| | KI-N Reverse | ACCACACAGGTGACTGAGGTA |
| | KI-C Forward | CTCAGGTGATGGAGTTCTGGCA |
| | KI-C Reverse | CTAAGGTGCAAGTCTCTCATAGAGCC |
| | *rp49* Forward | GCTAAGCTGTCGCACAAATG |
| | *rp49* Reverse | GTTCGATCCGTAACCGATGT |

**Supplementary Table 2. Antibodies for Western blotting (WB) and Immunohistochemistry (IHC) staining**

| Antibody | Source | Cat # | Dilution |
|---|---|---|---|
| Rabbit anti-human PNPO | Novus | NBP1-87302 | 1:500 (WB); 1:300 (IHC) |
| Mouse anti-beta tubulin | DSHB | E7 | 1:500 (WB) |
| Goat anti-rabbit HRP | Jackson ImmunoResearch | 111-035-144 | 1:10,000 (WB) |
| Goat anti-mouse HRP | Jackson ImmunoResearch | 115-035-003 | 1:10,000 (WB) |
| Mouse anti-Brp | DSHB | nc-82 | 1:10 (IHC) |
| Flurophore conjugated Donkey anti-primary antibody species | Life Technologies | A21206; A21202; A31573; A31571 | 1:400 (IHC) |
| Flurophore conjugated Donkey anti-primary antibody species | Jackson ImmunoResearch | 715-585-150; 711-585-152 | 1:400 (IHC) |

#### References

[1] A.W.K. Frankel and Jr. G.E.Brosseau. A drosophila medium that does not require dried yeast. *Drosophila. Inf. Service*, 43:184, 1968.

[2] Kei Ito, Kazunori Shinomiya, Masayoshi Ito, J. Douglas Armstrong, George Boyan, Volker Hartenstein, Steffen Harzsch, Martin Heisenberg, Uwe Homberg, Arnim Jenett, Haig Keshishian, Linda L. Restifo, Wolfgang Rössler, Julie H. Simpson, Nicholas J. Strausfeld, Roland Strauss, and Leslie B. Vosshall. A systematic nomenclature for the insect brain. *Neuron*, 81:755–765, 2 2014.
